# Transcription factor binding site divergence across maize inbred lines drives transcriptional and phenotypic variation

**DOI:** 10.1101/2024.05.31.596834

**Authors:** Mary Galli, Zongliang Chen, Tara Ghandour, Amina Chaudhry, Jason Gregory, Miaomiao Li, Xuan Zhang, Yinxin Dong, Gaoyuan Song, Justin W. Walley, George Chuck, Clinton Whipple, Heidi F. Kaeppler, Shao-shan Carol Huang, Andrea Gallavotti

## Abstract

Regulatory elements are important constituents of plant genomes that have shaped ancient and modern crops. Their identification, function, and diversity in crop genomes however are poorly characterized, thus limiting our ability to harness their power for further agricultural advances using induced or natural variation. Here, we use DNA affinity purification-sequencing (DAP-seq) to map transcription factor (TF) binding events for 200 maize TFs belonging to 30 distinct families and heterodimer pairs in two distinct inbred lines historically used for maize hybrid plant production, providing empirical binding site annotation for 5.3% of the maize genome. TF binding site comparison in B73 and Mo17 inbreds reveals widespread differences, driven largely by structural variation, that correlate with gene expression changes. TF binding site presence-absence variation helps clarify complex QTL such as *vgt1*, an important determinant of maize flowering time, and DICE, a distal enhancer involved in herbivore resistance. Modification of TF binding regions via CRISPR-Cas9 mediated editing alters target gene expression and phenotype. Our functional catalog of maize TF binding events enables collective and comparative TF binding analysis, and highlights its value for agricultural improvement.

## Introduction

TFs control when and where genes are expressed (Kim and Wysocka 2023). As a result, loss-of-function TF mutations can result in dramatic phenotypic changes resulting from widespread perturbation of downstream transcriptional networks. In contrast, subtle phenotypic changes can be achieved by modifications to non-coding, *cis*-regulatory regions, which can quantitatively alter TF target gene expression (e.g. pattern, level, condition) (Rodriguez-Leal et al. 2017). In plants, the application of knowledge about these TFs, their binding sites, and the features necessary to modulate gene expression and thereby phenotype is unique relative to animals where the main focus is on their correlation with disease phenotypes to improve human health (Weirauch et al. 2014) (Barrera et al. 2016). Instead, in plants, trait variation and engineered phenotypes hold the promise to allow for the generation of new varieties that address pressing food security needs (Wallace, Rodgers-Melnick, and Buckler 2018). These include crops that are resilient to extreme weather such as drought and flooding, varieties that harbor changes in vegetative and inflorescence architecture enabling higher yields, and/or plants that show adaptation to growth in day length sensitive regions or artificial settings such as vertical farming.

It has been estimated that a large percentage of phenotypic diversity maps to non-coding regions (Rodgers-Melnick et al. 2016) (Spielmann and Mundlos 2016) (Chen et al. 2024) (Maurano et al. 2012) (Springer, de Leon, and Grotewold 2019) (Aguirre et al. 2023; Ricci et al. 2019). Because these regions are small and tend to be less conserved than coding regions, identification of functional non-coding spaces is challenging. Maize is known for having high phenotypic and nucleotide diversity, although the degree to which TF binding differences contribute to these factors is unknown. Here we empirically mapped TF binding for over 200 TFs in two distinct maize inbred lines. We found many conserved TF binding events across the two genomes that when incorporated with orthogonal regulatory information such as unmethylated regions (UMRs), accessible chromatin (ATAC-seq and MOA-seq), and histone modification data pinpointed both proximal and distal functional regulatory regions with single bp accuracy. We also found evidence for extensive TF binding variation across the two genomes, identifying genotype-specific TF binding events and positional variants in which TF binding events relocate over 10kb or greater distances. These differences were often associated with changes in target gene expression across the genotypes, providing direct evidence for how TF binding diversity shapes phenotypes. Furthermore, using CRISPR-based editing of TF binding sites, we show that modifications to gene expression and phenotypes can be rationally engineered.

## Results

### Large scale mapping of TF binding sites in maize B73

The maize genome contains over 2,500 DNA-binding TFs comprising over 40 families, with some families containing only a few members while others contain hundreds (Burdo et al. 2014; Blanc-Mathieu et al. 2024). Relatively few of these TFs have been characterized genetically in maize, although those that have are known to control critical processes with agronomic importance suggesting that further functional characterization of additional TFs will be advantageous (Richardson and Hake 2022). While classical genetic studies have focused on the characterization of single TFs or those from particular biological processes, there is a need to understand how plant TFs work in an integrated manner to control gene expression. To broadly assess DNA-binding properties across many families in an unbiased manner and thereby maximize diversity, we selected a subset of TFs from various DNA-binding TF families for DAP-seq experiments using genomic DNA from the maize reference genome B73. Specifically, we tested 300 unique TFs and obtained high quality data for 166 maize TFs. Combined with data from previous studies (Galli et al. 2018) (Ricci et al. 2019) (Dong et al. 2020) (Dai et al. 2022) (Wu et al. 2023) (Bang et al., 2024), this provided DAP-seq data for 30 unique DNA-binding TF families out of 36 total families tested (Supplemental Figure 1a). The number of peaks identified for each TF ranged from 1,153 to 122,695 giving an average of ∼33,000 peaks (Supplemental Figure 1b, Supplemental Table 1). Due to the *in vitro* nature of DAP-seq, these peaks represent all potential binding events regardless of tissue type or condition (O’Malley et al. 2016) (Zhang et al. 2021). Overall, 1.6 million non-redundant peaks were identified, with their collective binding sites corresponding to 5.3% of the maize genome. Putative target genes were assigned to each peak that resided within 10kb of the gene transcription start site (TSS), producing on average 8,400 target genes per TF dataset (Supplemental Figure 1c). Together these datasets provided a comprehensive view of DNA binding across the B73 maize genome, identifying exact chromosome coordinates for thousands of binding events in putative target genes and the motifs located within. Due to the large size of the maize genome and the frequent insertion of transposons near genic regions that often relocate regulatory elements far from TSSs (Lu et al. 2019; Ricci et al. 2019), this information is crucial for identifying gene regulatory elements and enhancers that impact gene function.

High resolution pairwise correlation analysis of DAP-seq derived TF binding events showed that similar TF family members bound similar sites that were largely distinct from those of other TF families, confirming the reproducible nature of the datasets and suggesting a high degree of intra-family similarity (Figure 1a). For example, many BZIP family members bound similar sites that were unique relative to those bound by members of the EREB, NAC or WRKY families, reflecting the stringent motif specificity conferred by the structurally unique DNA-binding domains of these different TF families (Figure 1a).

**Figure 1.**
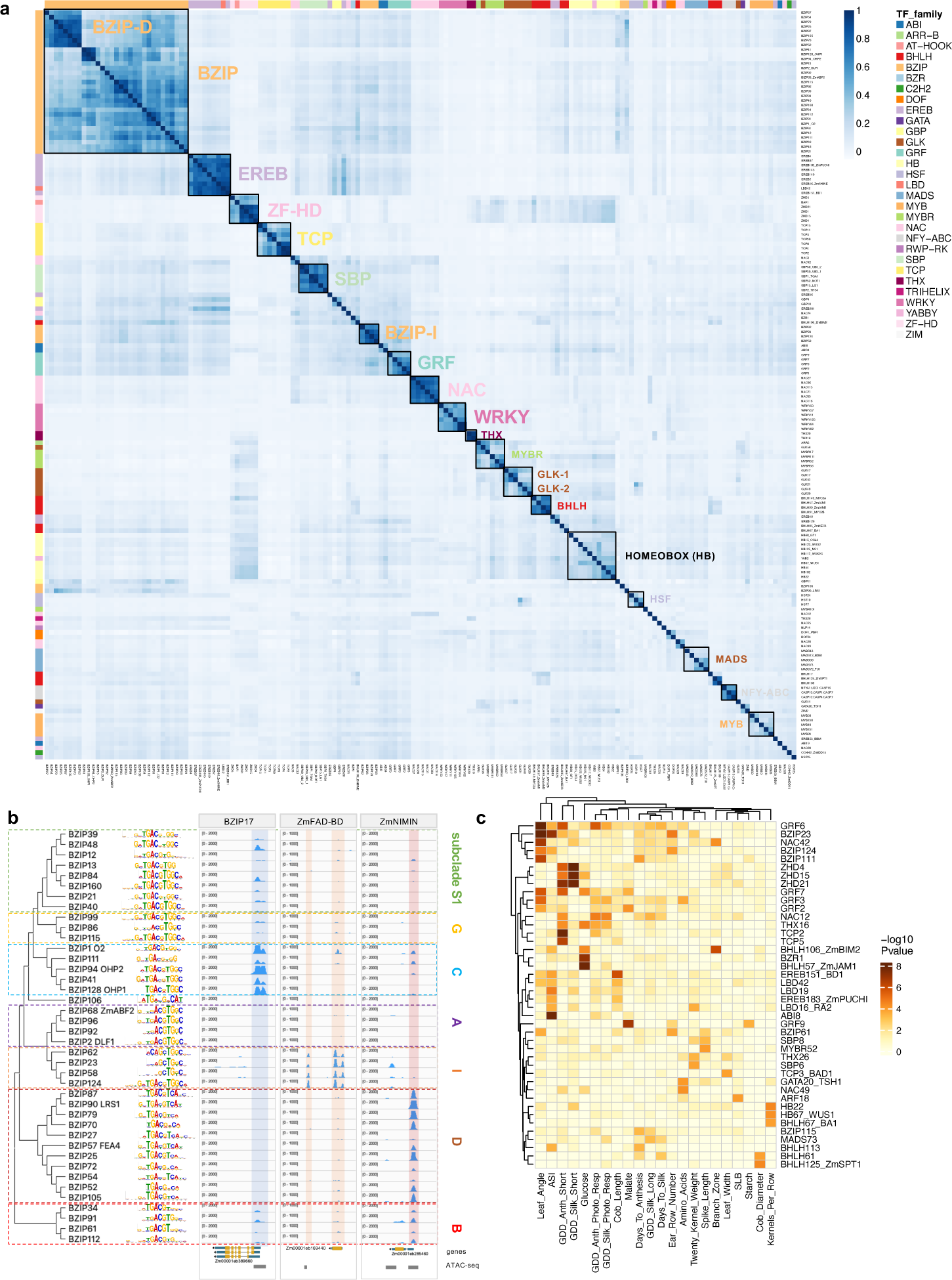
Large scale DAP-seq profiling of maize TFs provides high quality genome-wide binding site data for B73v5. **a.** Heatmap showing Pearson correlation of TF binding sites genome wide (10 bp bins). Side annotation colors correspond to TF structural families. **b.** Phylogenetic tree showing sub-clade specificity among sequence motifs and target sites of maize BZIP proteins. Loci shown include known targets of maize BZIPs or their Arabidopsis homologs. **c.** Heatmap showing TFs whose binding sites were enriched for overlap with published maize GWAS hits for various traits.

In several instances, sub-clade specific binding preferences were observed among members belonging to the same structural TF family. For example, BZIP sub-clades I and D showed low binding site correlation (Figure 1a). These genome-wide differences could also be seen at the individual locus level where certain BZIP sub-clade members showed binding in the putative regulatory regions of known target genes or maize homologs of known or inferred Arabidopsis TF-target gene pairs. For example, BZIP17 is a known target of sub-clade C member BZIP1/O2 (Zhan et al. 2018), while ZmFAD-BD, a maize homolog of AtFAD-BD previously shown to be bound by sub-clade I member AtBZIP59 (Xu et al. 2018) was bound by multiple maize sub-clade I members (Figure 1b). Additionally, ZmNIMIN known to be involved in salicylic acid signaling and regulated by the sub-clade D BZIPs (Hermann et al. 2013) was bound by several maize sub-clade D members (Figure 1b). In many cases, these sub-clade-specific binding site locations were also reflected as differences in motif preference, although not always, suggesting that sequences outside of the core binding motif influence binding site selection (Figure 1b). Similar observations were made for the LBD family (class I and II (Majer and Hochholdinger 2011)), EREBs (PLT/AIL, BBM, RAV, DREB), MYBs, NACs, and TCPs (Supplemental Table1). Phylogenies with corresponding motifs showed similar patterns in both maize and Arabidopsis (Supplemental Figure 2, 3). In addition to showing unique binding site preferences, certain TFs and TF families were largely enriched in unique GO biological functions that supported known roles (i.e. HSF24 targets were strongly enriched in heat stress related categories) and assigned putative novel functions to certain uncharacterized TFs, such as a role for NAC74 in immune responses (Supplemental Figure1d).

To further support the biological relevance of our TF binding site data, we assessed the overlap of peaks from our DAP-seq TFs with GWAS hits from 41 measured traits (including those related to plant architecture, development, disease resistance, and various metabolites) segregating within 5,000 recombinant inbred lines of the NAM population that captures much of the global diversity of cultivated maize (Wallace et al. 2014). Significant enrichments were found for twenty-two traits between the trait-associated SNPs and DAP-seq peaks of at least one TF (P<=0.001; Figure 1c). These included several GRFs and BZIPs associated with leaf angle (important for light capture) (Peng et al. 2021), BZIPs and ERFs associated with anthesis-silking interval (ASI; an important flowering time trait), SBP6 associated with kernel weight, and TCP3/BAD1/WAB associated with leaf width (Figure 1c). Conversely, for several well-characterized developmental TFs we observed enrichment of their DAP-seq peaks overlapping with GWAS hits for traits relevant for the TF function, such as architecture-related traits for SBP30/UB3 and flowering time-related traits for MADS73 (Supplementary Figure 4a, 4b). These observations held true across several GWAS significance thresholds, supporting the robustness of the results (Supplementary Figure 4a, 4b). Collectively, these data reveal the high quality and functional relevance of our large-scale genome-wide profiling of maize TF families.

### Impact of TF heterodimerization on DNA-binding and binding site preference

Protein-protein interaction screens have revealed that many TFs heterodimerize with either related family members and/or partner TFs from other families. These interactions have the potential to alter binding site specificity and thus expand the repertoire of TF binding sites beyond those achieved by single TFs. However, the impact of protein interactions on DNA binding in plants has only been investigated for a few families at the genome-wide level, despite its importance and implications for crop improvement (Lai et al. 2020) (Li et al. 2023). For example, breeding-based selection for certain traits often acts upon quantitative changes in TF expression levels (Alonge et al. 2020) (Aguirre et al. 2023); for dosage sensitive TFs whose interactions with partners influences DNA binding, such selections would be best informed by knowledge of protein partners and how those protein interactions affect DNA binding.

Among DNA-binding TF families, several employ homodimeric and/or heterodimeric DNA binding domains (DBDs), in which a functional DBD is formed by the dimerization of two identical or distinct monomers. To explore further how combinatorial heterodimeric interactions among TF family members impact DNA binding specificity we employed doubleDAP-seq, a modified version of DAP-seq for the analysis of heterodimeric complexes (Li et al. 2023) (Supplemental figure 5a). We focused on the large maize BHLH family which has 175 members comprising twelve main groups based on structural similarities to Arabidopsis BHLHs (Heim et al. 2003). The BHLH DNA-binding domain is highly conserved and consists of an alpha helix containing several basic residues that interact with DNA, followed by a loop and a second alpha helix that mediates the formation of BHLH dimers. The ‘basic’ region of each monomer contacts a half site of the E-box (CANNTG) or the canonical palindromic E-box variant CACGTG (G-BOX). Among the twelve main plant BHLH groups, nearly all contain highly conserved H-E-R amino acid residues in their ‘basic’ region which have been shown to contact DNA (Heim et al. 2003) (de Martin, Sodaei, and Santpere 2021). One exception is the group VIII BHLHs which include the Arabidopsis *HECATE* (*HEC*) and *INDEHISCENT* (*IND*) genes involved in meristem and female reproductive organ development (Gaillochet et al. 2017) (Crawford and Yanofsky 2011) and the maize *BARREN STALK1* (*BA1*) gene involved in inflorescence architecture (Gallavotti et al. 2004). A total of nineteen members comprises this group in maize (Supplemental Figure 5b), with all containing highly conserved Q-A-R residues in place of the H-E-R residues (Figure 2a), leading to speculation that they may not directly bind DNA. Alpha-fold based structural prediction (Jumper et al. 2021) (Mirdita et al. 2022) however revealed that this domain forms an alpha helix resembling that of the canonical ‘basic’ domain in other BHLHs, albeit one that is shorter by two helical turns relative to canonical BHLH regions (Figure 2b). We therefore first used the traditional DAP-seq assay to test the capacity of group VIII members to bind DNA. Of the nine members tested, eight yielded a moderate number of peaks (1153 to 19,154 peaks) that were enriched for a similar, but non-canonical motif highly divergent from the E-box (Figure 2c). These findings suggest that certain Q-A-R type BHLHs are capable of binding DNA despite lacking non-canonical DNA-contacting amino acid residues in their basic regions.

**Figure 2.**
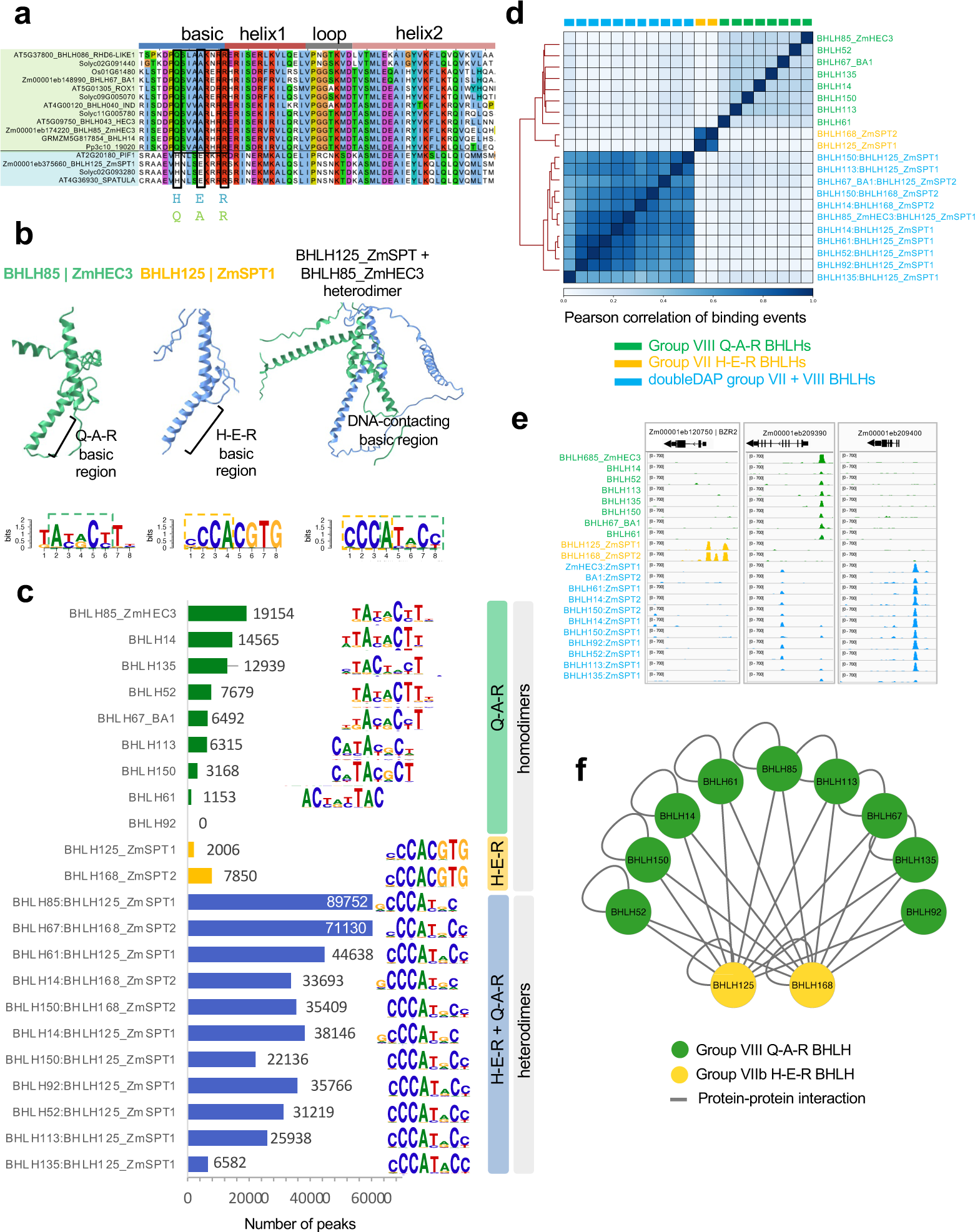
DoubleDAP-seq analysis of Q-A-R type BHLHs. **a.** Clustal alignment of amino acids in BHLH DNA binding domain of Q-A-R BHLHs (light green; group VIII) and group VIIb BHLHs (light blue) with QAR and HER residues shown. **b**. AlphaFold predicted ribbon diagrams of BHLH85 (Q-A-R), BHLH125 (H-E-R), and the heterodimer. Empirically determined DAP-seq motifs are shown below each structure. **c**. Number of peaks and top motifs called for DAP-seq and doubleDAP-seq datasets. **d**. Pearson correlation of genome-wide binding events. **e.** Genome browser screenshots of binding by homo- and heterodimers. **f.** Summary of protein-protein interactions based on DAP-seq experiments.

Previous studies have indicated that Arabidopsis HEC and IND (clade VIII BHLHs) also physically interact with SPATULA (SPT) and ALCATRAZ (ALC) (Gremski, Ditta, and Yanofsky 2007) (Girin et al. 2011), two BHLHs that belong to subclade VIIb and contain the canonical H-E-R basic residues. Maize contains two orthologs of SPT/ALC (BHLH125/ZmSPT1 and BHLH165/ZmSPT2), both of which are expressed in multiple tissues throughout plant development (Supplemental Figures 5b and 6a), and when tested in the single protein DAP-seq assay, bound a moderate number of peaks that were enriched for the canonical E-box motif CACGTG (Figure 2b). Surprisingly, testing these BHLHs in doubleDAP-seq revealed that heterodimers formed by ZmSPTs and the clade VIII Q-A-R members yielded peak numbers up to 44 times higher than those formed by Q-A-R sub-clade VIII homodimers (Figure 2b, 2c, 2f). Furthermore, a highly enriched, unique motif (CCCATnCC) was identified in sites bound by the cross-clade heterodimers (Figure 2b). This motif differed substantially relative to that which was enriched in either the Q-A-R or SPT-type homodimers (Figure 2c), and consequently resulted in distinct binding locations for all three BHLH dimer combinations (Figure 2d, 2e). The CCAT half-site bound by the heterodimer resembled the CCAG half-site bound by ZmSPTs, with the major change being the preference of G instead of T in the ‘N’ position of the known BHLH ‘CAN’ half site (De Masi et al. 2011), while the other half site resembled that of BHLH67/BA1 and BHLH85/ZmHEC3 (Figure 2b, c). Our DAP-seq approach has thus demonstrated that the plant-specific Q-A-R type DNA binding domain binds DNA both as a homodimer and a heterodimer with SPT-related BHLHs, thereby expanding the known BHLH binding repertoire and potentially explaining the dual roles that have been proposed for *HEC* genes (Gaillochet et al. 2017). In many cases, we noted that new binding events of the ZmSPT-QAR heterodimers shifted from their proximal location near TSSs to distal intergenic regions (Supplemental Figure 6b), suggesting one possible role for group VIII HEC genes could be to sequester ZmSPTs from their homodimeric target sites.

Taken together our doubleDAP-seq results revealed that certain sub-clades of BHLHs form heterodimers that alter DNA binding site specificity and offer a convenient method to test for individual TF contributions to DNA binding. These experiments expanded our understanding of the TF binding repertoire, identifying a novel, plant specific BHLH binding motif bound by Q-A-R type group VIII family members and their heterodimeric partners.

### Integration of TF binding events reveals cooperative regulatory potential

Because of the high degree of intra-family binding site overlap observed for the TFs analyzed in this study and previous DAP-seq analysis (Galli et al. 2018), we condensed our datasets into a panel of 66 TFs, (hereafter ‘maize TF diversity panel’, Supplemental Table 2), designed to focus classification efforts for further analysis. Only datasets that showed unique binding motifs, belonged to a distinct subfamily, and/or had a Pearson correlation less than 0.5 were included (Figure 3a; Supplemental Figure 7a). We note that TFs excluded from this list however should not necessarily be considered redundant as many exhibited several unique binding sites and/or showed unique expression patterns, suggesting they play a non-redundant role in maize development. For example, while many SBP TFs showed similar motifs and had highly correlated genome-wide binding profiles, their individual RNA expression patterns often showed tissue specificity which could be a large driver of non-redundant functionality (Supplemental Figure 7b, 7c). For this reason, we recommend that TFs in the diversity panel should be interpreted as representative members of the TF family or sub-family, and any follow-up investigation should consider tissue specificity and the possible role of closely related family members.

**Figure 3.**
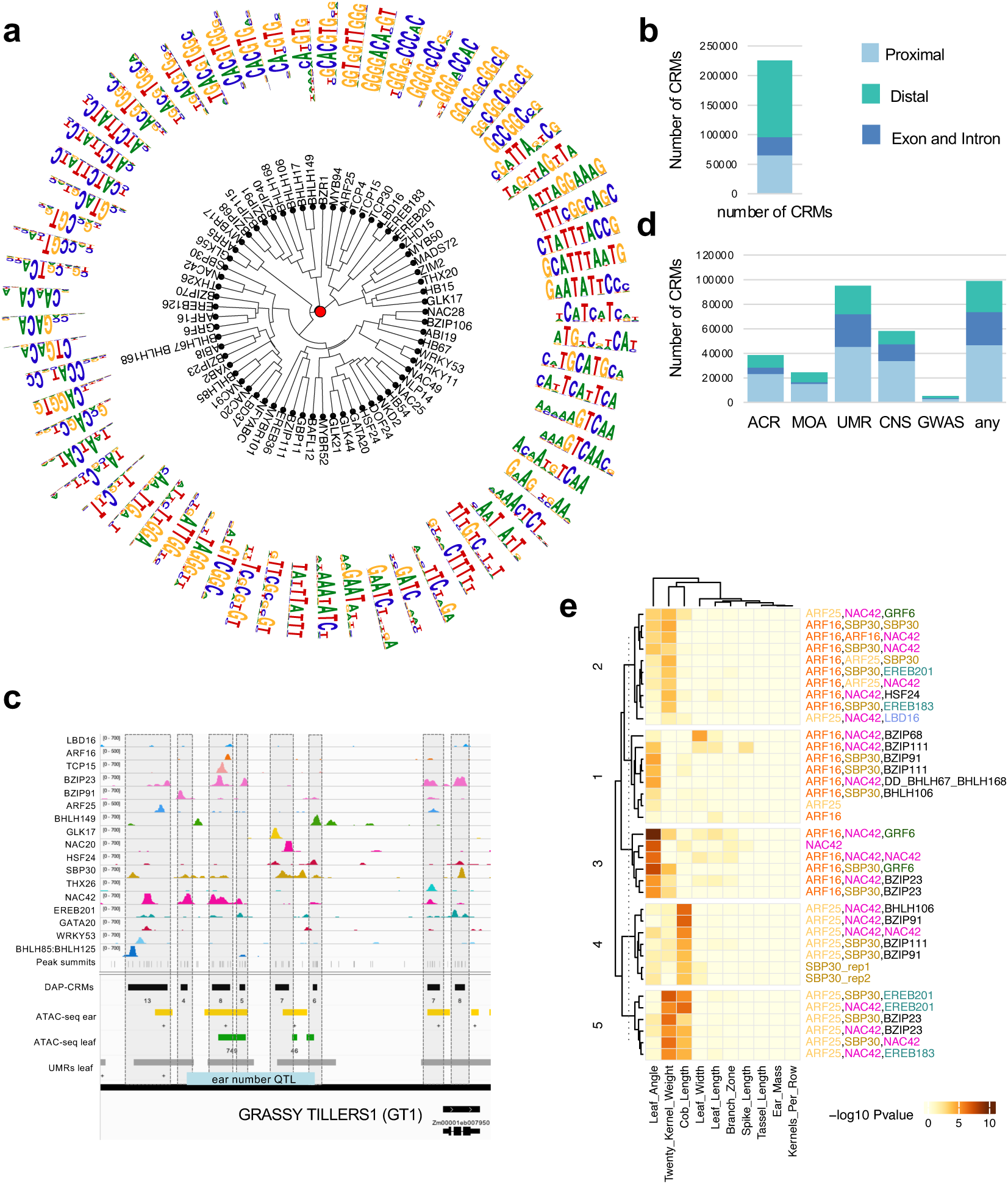
Maize TF diversity panel reveals functional *cis*-regulatory modules. **a**. Maize TF diversity panel is represented by mostly distinct motifs. **b**. Stacked bar graph showing the number of DAP-CRMs (clusters of TF binding sites) and their prevalence in various gene features. **c**. Genome browser screenshot of TF binding peaks within the GRASSY TILLERS1 locus which contains eight distinct DAP-CRMs, each with three or more TFs. The estimated region of the known prolificacy QTL is also shown. **d**. Overlap of DAP-CRMs with various orthogonal functional datasets. **e.** Heatmap showing DAP-CRMs with specific combinations of TFs that are enriched for different GWAS traits at a significance threshold of 1e-7.

Open chromatin profiling methods such as ATAC-seq have identified many regions of functional non-coding space, however empirical evidence of which TFs bind in these regions is needed on a large scale. We therefore examined the overlap of TFs from our diversity panel with orthogonal functional regulatory features including regions of accessible chromatin (ATAC-seq and MOA-seq), unmethylated regions (UMRs; tissue/condition agnostic markers of regulatory regions), histone modification marks, conserved non-coding sequences from sorghum, and published leaf protoplast ChIP-seq data (Savadel et al. 2021) (Crisp et al. 2020; Song et al. 2021; Ricci et al. 2019; Tu et al. 2020; Hufford et al. 2021). We observed a variable degree of overlap depending on the TF (Supplemental Figure 7d), suggesting diverse transcriptional roles. These data help further define our understanding of regulatory regions at the TFBS level and guide GWAS analysis.

The large number of TFs profiled in our DAP-seq experiments allowed for parallel analysis of binding events for multiple TFs from various families at individual loci. As has been noted in other plant and animal studies, we observed that maize TFs often bound in clusters, forming distinct *cis*-regulatory islands whose collective role is still unclarified (Marand et al. 2023) (Schmitz, Grotewold, and Stam 2022) (Tu et al. 2020) (Jores et al. 2024). To better characterize these clusters, we collapsed all peak summits from our TF diversity panel that were within 300bp of each other and selected those containing three or more TF binding events, identifying 225,235 DAP-*cis*-regulatory modules (DAP-CRMs) to which each was assigned a unique tracking identifier (Figure 3b). DAP-CRMs contained a mean of 5.3 TFs and were on average 344bp with a maximum length of 4.0kb (Supplementary Figure 8a,b). In total, they covered 3.6% of the maize genome. Twenty-nine percent resided within 10kb of a gene, 14% were in the gene body, and 57% were classified as distal DAP-CRMs (defined as >10kb upstream from the TSS or >300bp downstream of the TES; Figure 3b, Supplemental Figure 8c). Among the several distal DAP-CRMs identified in our analysis, several overlapped genetically defined QTL such as *GRASSY TILLERS* and *vgt1* providing information on which TFs may be controlling these genes (Figure 3c, Supplemental Figure 9a), (Wills et al. 2013) (Salvi et al. 2007). This approach also allowed us to define novel distal regulatory regions and TF binding sites within that likely control characterized genes such as *UNBRANCHED2* (*UB2*) and *INDETERMINATE GAMETOPHYTE* (*IG1*) for which little is known at the regulatory level (Supplemental figure 9b, 9c) (Chuck et al. 2014) (Evans 2007).

To build upon previous efforts to map functional regulatory regions genome-wide, we next incorporated the DAP-CRMs with existing orthogonal datasets. Overlaying our DAP-CRMs on accessible chromatin region (ACR) data from leaf and ear ATAC-seq (Hufford et al. 2021) (Ricci et al. 2019) and leaf unmethylated regions (UMRs; tissue/condition agnostic markers of functional regulatory regions, (Crisp et al. 2020) revealed that about 17% overlapped with ACRs, while 42% overlapped with UMRs, a six and four-fold respective enrichment relative to randomly shuffled coordinates (Figure 3d). Similarly, 26% of DAP-CRMs overlapped with conserved non-coding sequences (CNS) from sorghum, a four-fold enrichment relative to randomly shuffled coordinates (Song et al. 2021). In addition, we were able to identify over 4,500 DAP-CRMs (∼2%) that overlapped GWAS SNPs (Wallace et al. 2014).

We next sought to understand if certain combinations of TFs were consistently localized together within our DAP-CRMs. To this end, we selected DAP-CRMs that overlapped GWAS loci related to vegetative and inflorescence architecture, and contained at least one peak for ARF, SBP, or NAC, TFs known for their roles in auxin biology and plant architectural traits. This analysis showed that in certain cases co-binding by specific TF combinations was more influential than individual TFs alone. For example, while ARF16 (activator), ARF25 (repressor), and SBP30/UB3 binding sites individually had minor enrichment for GWAS hits, the DAP-CRMs containing combinations of ARFs, SBPs, and NACs were much more enriched than the individual TFs (Figure 3e). Furthermore, different combinations of TFs showed enrichment for different traits. For example, ARF16-containing DAP-CRMs that also had SBP30 and NAC42 sites were highly enriched for SNPs associated with leaf angle, while ARF25-containing DAP-CRMs that had SBP30 and NAC42 sites were instead highly enriched for cob length (Figure 3e). Similarly, ARF25- and EREB-containing DAP-CRMs were specifically enriched for kernel weight. These data support a model where clusters of specific sets of TFs influence specific traits.

Overall, the DAP-CRMs identified here revealed functional composite regulatory regions as well as the TF families that bind them, either alone or as potential cooperative units. Such TF clustering may allow integration of multiple cellular signals to regulate gene networks, act in distinct cell types, and/or control specific aspects of pleiotropic phenotypes.

### Comparative TF binding in two diverse inbred lines

Maize was domesticated from its wild progenitor teosinte around 10,000 years ago, and intensive selection and modern breeding in the past 100 years have resulted in the generation of highly diverse inbred lines which can be grown in regions far from its original tropical location. The release of several maize genome assemblies has revealed that a core set of genes are conserved in related accessions (Hufford et al. 2021). However, such figures have not yet been empirically explored for TF binding sites despite the general belief that differences in individual TF binding sites could be significant drivers of phenotypic differences among inbreds (Rodgers-Melnick et al. 2016) (Yocca and Edger 2022). Similarly, little is known about the composition and flexibility of TF binding sites even within conserved *cis*-regulatory regions that likely correspond to crucial regulatory regions dictating core gene expression programs. Identification and characterization of these regions would allow the construction of a pan-genomic TF binding regulatory space that could assist breeding efforts.

Two inbred lines that have been widely used in maize breeding are Mo17 and B73 (Sun et al. 2018). Important phenotypic differences between the two genotypes include cold tolerance, disease susceptibility, flowering time, and plant architecture (Eichten et al. 2011). Whole genome assemblies are available for both genomes, and it has been reported that they are replete with large structural variations (SVs; defined here as indels >50bp), SNPs, and indels (<50bp) (Sun et al. 2018). Given that each of these features overlapped with a sizeable percentage of the peaks identified in our B73 DAP-seq datasets (Supplemental Figure 10a), we hypothesized they were likely to contribute to TF binding site differences. Therefore, to better understand empirically how sequence features influence TF binding and in general investigate how TF binding sites varied across different maize inbreds, we performed DAP-seq experiments using Mo17 genomic DNA and the 200 maize TFs that gave positive peak results in B73 either in this study or previous DAP-seq studies. As with the B73 DAP-seq experiments, direct mapping of Mo17 DAP-seq reads to the Mo17 genome identified a range of peaks (Supplemental Table S1). Importantly, Mo17 datasets reported nearly identical motif enrichment results to those obtained in B73 (Supplemental Table S1), supporting the highly reproducible nature of our DAP-seq approach (O’Malley et al. 2016).

We next sought to directly quantify how TF binding peaks varied among the two genomes. Using TFs from our TF diversity panel, we mapped reads to their tested genome, equalized read numbers across the datasets, and then performed coordinate ‘liftover’ using a high coverage whole genome syntenic alignment generated by Anchorwave (Song et al. 2022) (Zhao et al. 2014). This approach converted genomic coordinates of Mo17 to those of B73 (and vice versa), allowing direct comparison of peak dataset coordinates. Whole genome Pearson correlation of read location for direct and lifted datasets showed strong genome wide correlation among matching TF datasets (Supplemental Figure 10b). Furthermore, very few non-cross-mappable peaks overlapped with non-aligned regions suggesting that our approach was likely capturing a large percentage of possible functional variation (Supplemental Figure 10c). Overall, comparison of peak coordinates for diversity panel TFs revealed that between 7-72% of peaks were specific to B73, while 28-93% of peaks were shared between both genomes (average 64% shared in B73 and Mo17; Figure 4a). Similar values were found for Mo17-specific peaks (Supplemental Figure 11a). TFs that showed the highest degree of shared peaks were those that had a high percentage of exonic binding (i.e. EREB183), while those with low percentages bound more frequently to distal intergenic regions (i.e. MYBR52). These values were conservatively estimated based on the top 20% of peaks of each TF to account for differences in peak calling at lower thresholds and did not consider quantitative differences in peak binding. The degree of shared TF binding sites was on average lower than that reported for ACRs (ATAC-seq and MOA-seq), UMRs, and coding regions (Noshay et al. 2021) (Hufford et al. 2021) (Sun et al. 2018) (Engelhorn et al. 2023) indicating TF binding is relatively more variable and less constrained.

**Figure 4.**
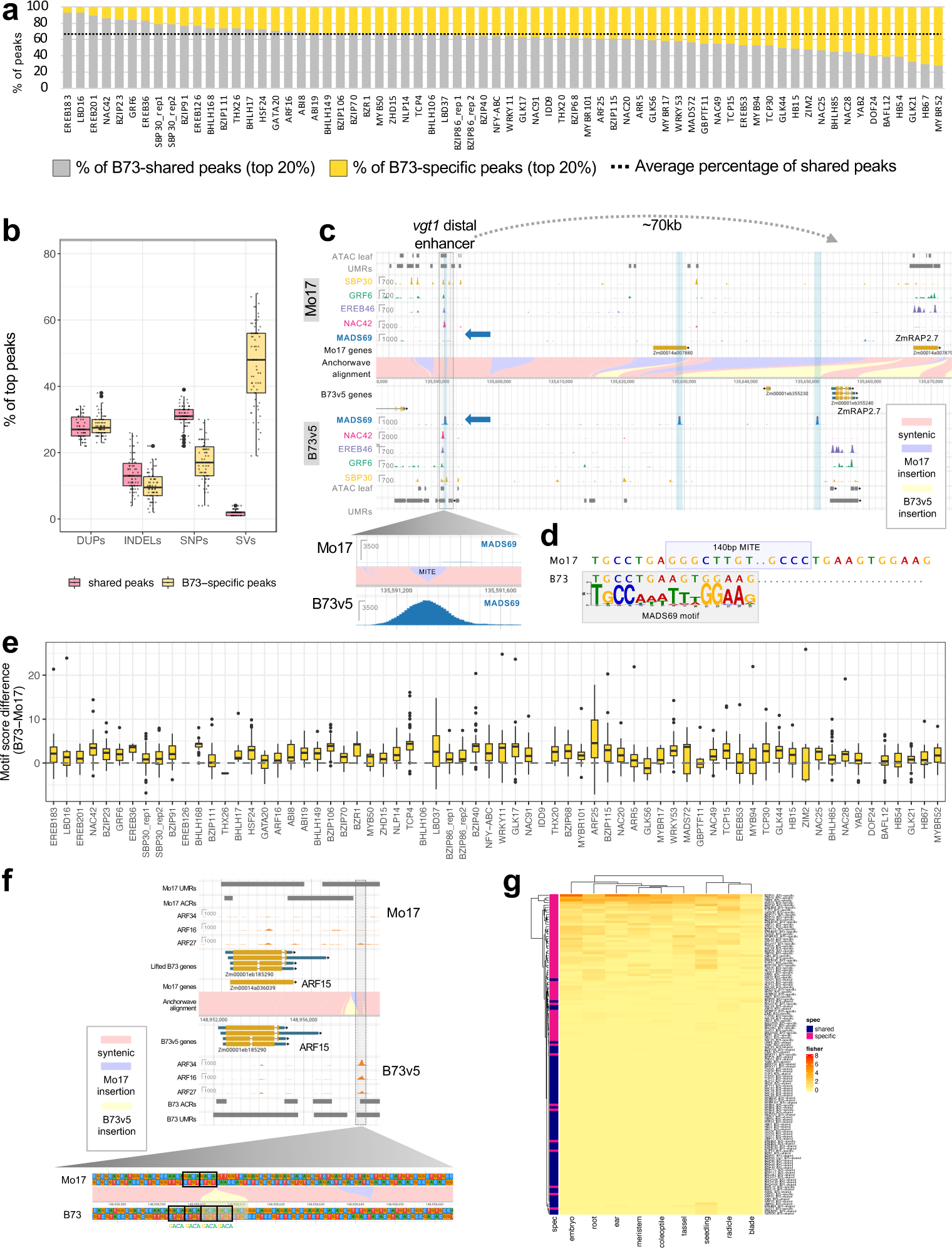
Genotype-specific peaks are prevalent in B73 and Mo17 and explain genetically defined QTL. **a**. Stacked bar graph showing the percentage of B73-specific peaks (yellow) and shared peaks found in both B73 and Mo17 (gray). Dotted line indicates the average percentage of shared peaks. **b**. Percentage of B73-specific (light yellow) and shared (pink) peaks that overlap with duplicated regions (DUPs), small indels less than 50bp (INDELs), SNPs, and structural variants (SVs). Each datapoint corresponds to the percentage of peaks from an individual TF overlapping the indicated category. **c**. JBrowse 2 screenshot of *vgt1-RAP2.7* locus showing three MADS69 binding events upstream of *RAP2.7* in B73v5, one of which is located in the genetically defined *vgt1* enhancer. Lower left panel shows a close-up of the region in Mo17 containing the MITE transposon and the corresponding region in B73v5 showing the MADS69 peak. **d**. Alignment of the MADS69 CArG-box motif with B73 and Mo17 sequences. The MITE inserts in the middle of the motif, eliminating high information content nucleotides within the motif. **e**. Box plots showing motif score differences for B73-specific peaks of each of the TFs in the diversity panel. For most TFs, B73 sequences have higher motif scores than the corresponding sequences in Mo17 within B73-specific peaks regions. **f**. JBrowse 2 genome browser screenshot showing example of 12bp indel that results in ARF TF binding in B73 but not Mo17 at the ARF15 locus. **g.** Heatmap showing enrichment scores of differentially expressed putative target genes of shared and B73-specific peaks in B73 vs. Mo17 in various tissues (Zhou et al., 2019). B73-specific target genes are indicated with a magenta bar and shared target genes are indicated with a dark blue bar. Heatmap color scale maps to -log 10 transformed FDR-adjusted P value of Fisher’s exact test.

We next measured the degree to which SNPs, indels (<50bp), and SVs (>50bp) impacted TF binding. In general, we noted a broad distribution of peak overlap with SNPs and indels for both shared and B73-specific peaks. On the other hand, B73-specific peaks showed a much higher percentage of their peaks that overlapped with structural variants relative to shared peaks (Figure 4b). These findings suggest that SVs are the biggest contributor to TF binding site variation across genotypes and could serve as major drivers of phenotypic variation. For example, we noted three large MADS69 peaks located upstream of RAP2.7 in B73 that were absent in Mo17. Two of the MADS69 peaks were located within a large B73-specific insertion proximal to RAP2.7, while a third was located ∼70kb upstream in the known *vgt1* enhancer region in B73 (Figure 4c). Strikingly, the absence of the *vgt1* MADS69 peak in Mo17 appeared to be caused by the insertion of a 140bp MITE transposon that has previously been strongly linked with early flowering time in Mo17 and other inbreds, but for which the underlying molecular cause has remained elusive (Buckler et al. 2009) (Salvi et al. 2007) (Castelletti et al. 2014). Our data indicate that a CArG-box binding site of MADS69 in B73, is bisected in Mo17 by the MITE, but that additional TF binding events within the *vgt1* enhancer remain intact (Figure 4c,d). Whether the absence of MADS69 binding in Mo17 is caused directly by disruption of the binding site or indirectly by methylation (Castelletti et al. 2014), remains to be determined. Furthermore, we note that MADS69 bound equally well at other loci in both B73 and Mo17, indicating the absence of binding in *vgt1* in Mo17 is not due to a lower quality Mo17 dataset (Supplemental Figure 11b). As both MADS69 and RAP2.7 are known regulators of flowering time in maize that are believed to be acting in the same pathway (Liang et al. 2019), our data could pinpoint the underlying molecular details of an important flowering time haplotype in maize. We also note that an additional flowering-time-related MADS member in our collection, MADS72/TUNICATE, also showed weak binding that overlapped the strong MADS69 peak in *vgt1*, suggesting that a complex interplay of MADS binding and/or multimeric interactions could occur *in vivo* at this site as is common with MADS-box genes (Smaczniak et al. 2017) (Hugouvieux and Zubieta 2018) (Supplemental Figure 9a).

While structural variations were the largest driver of inbred-specific peaks, we also observed SNPs and INDELs that contributed to binding site variation. To explore the extent to which SNPs affected TF binding, we computed the differences in motif scores between the B73-specific peak sequences and their syntenic Mo17 sequence regions, revealing that B73-specific peak sequences were more likely to have higher TF motif match scores than syntenic Mo17 sequences that did not have a peak (Figure 4e). Additionally, among the peaks that overlapped with SNPs, we found that B73-specific peaks had a higher percentage of peaks with four or more SNPs per peak compared to peaks with only a few SNPs (Supplemental Figure 11c). The opposite trend was seen for shared peaks, where a higher percentage of peaks had only a few SNPs per peak relative to those with many SNPs per peak. This suggests that TF binding is sensitive to SNPs residing in the sequence motif and the number of SNPs within a region, as expected. Small indels less than 50bp also affected TF binding, often in unexpected ways. For example, ARF peaks were observed downstream of ARF15 in B73 but were absent in Mo17 due to a 12bp insertion in B73 that resulted in four direct repeats of the core ARF binding site TGTC/GACA, producing an optimal TF binding site spacing pattern previously shown to be preferred by ARF- A members (Figure 4f) (Boer et al. 2014) (Galli et al. 2018). Overall, these data demonstrate the power of our comparative TF mapping approach to identify specific TF binding site footprints that likely contribute to phenotypic variation.

To further investigate if additional genotype-specific peaks were associated with previously identified functional regions, we calculated their overlap with orthogonal functional datasets (UMRs, ACRs, MOA-seq and CNS from sorghum). We found that B73-specific peaks showed between two-fold and six-fold higher overlap with orthogonal datasets compared to random shuffling of peak coordinates (p < 0.0001, unpaired Student’s t-test), similar to shared peaks which showed ∼four to eight-fold enrichment relative to shuffled coordinates (p < 0.0001, unpaired Student’s t-test; Supplemental Figure 11d), indicating that many B73-specific peaks were supported by additional functional data. These findings show that putative enhancers and their TF composition are often dynamic across genotypes.

While genotype-specific binding sites are likely drivers of phenotypic differences between genotypes, highly conserved TF binding sites and CRMs are likely to underlie core processes and developmental programs. We noted many instances of highly conserved TF binding sites and DAP-CRMs that had identical composition and spacing in both B73 and Mo17. For example, *MADS67*, a gene shown to increase yield when overexpressed in transgenic lines (Wu et al. 2019), showed binding by NLP14, EREB46, a NFY-ABC trimer, and BHLH57 in exactly the same location in both genotypes regardless of three SNPs that overlapped the BHLH57 site (Supplemental Figure 12a).

In contrast to the shared TF binding sites that appeared positionally constrained, we also observed many other shared TF binding sites that could be classified as positional variants (PosVs), in which TF binding events were present in both B73 and Mo17 but located at variable distances from the TSS of their common putative target gene. For example, ZmATL6, an E3 ubiquitin ligase whose Arabidopsis homolog is involved in defense and carbon/nitrogen response (Maekawa et al. 2012), showed proximal regulatory binding by BZIP111 and NAC42 in both B73 and Mo17, however additional conserved binding events (BZIP111, HSF24, NAC3, NAC42, SBP8, SBP30, and THX26) were relocated 11kb upstream in Mo17 due to a sequence insertion (Supplemental Figure 12b). Such positional information regarding the relative location of regulatory regions is important because it may i) quantitatively influence target gene expression, and ii) inform promoter construct generation and/or guide CRISPR-mediated editing of regulatory regions. Among the TFs in our TF diversity panel, we noted that ∼2-10% of their B73 peaks that resided in upstream regulatory regions (i.e. 5’UTR and - 10kb promoter) were relocated 500bp or more from their position relative to the TSS in Mo17 (average 5%; Supplemental Figure 12c). This percentage was substantially lower on average than peaks that were relocated less than 500bp from the TSS in both genotypes and those that were unique to a specific genotype, but could still be an important source of expression variation given that in many cases peaks relocated greater than 10kb from their presumptive target gene.

### Correlation of genotype-specific peaks with RNA expression

Having noted a substantial number of TF binding differences (both presence/absence and posVs), we next asked how many of these were associated with expression differences in putative target genes. We first called target genes for B73-specific peaks and shared peaks, using only those genes that contained peaks within 10kb upstream to 300bp downstream of the TSS. This data was then compared to differentially expressed genes between B73 and Mo17 in multiple tissues (FDR adjusted P value < 0.05, FC > 4) computed from a published RNA-seq dataset (Zhou et al. 2019). We found that genes associated with B73-specific peaks more significantly overlapped with differentially expressed genes between B73 and Mo17 compared to the genes associated with shared peaks, indicating that genotype-specific TF binding events can impact genotype-specific gene expression (Figure 4g). We also investigated how positional variation impacted gene expression. For many TFs, a significantly higher percentage of target genes were differentially expressed between B73 and Mo17 when their associated DAP-seq peaks were relocated more than 500bp, compared to those genes for which the associated DAP-seq peaks were relocated less than 500bp. This suggests that variation in binding site location could impact gene expression (Supplementary Figure 12d).

We were also interested in whether TF binding events were drivers of cell-type specificity. To investigate this, we used cell type specific root expression data (Guillotin et al. 2023) to examine the overlap with our B73-specific and B73-Mo17 shared target genes. Interestingly, we found *shared* target genes were more enriched than B73-specific target genes for many root cell type marker genes, indicating that TF-driven expression patterns that determine core cell types are conserved across genotypes (Supplemental Figure 12e). This data agrees with reports showing that TF-target gene pairs are also often conserved across species (Guillotin et al. 2023).

### Induced TF binding variation impacts gene expression and phenotype for important agronomic traits

To further validate the value of our DAP-CRM maps, we performed CRISPR-based genome editing of TF binding sites at several maize loci, including sites located in the upstream regulatory regions, a 3’ UTR, and distal enhancers. In most cases, deletion of TF binding sites led to altered expression and/or phenotypes and allowed us to pinpoint specific TFs affecting phenotypic output. For example, deletion of a DAP-CRM region containing at least nine TF binding events including an SBP30/UB3 and SBP2/TSH4 binding site in the upstream non-coding region of the *TSH1* gene led to a classic *tsh1* mutant phenotype that resembled loss-of-function mutations, with reduced tassel branching and extended outgrowth of bracts, typically suppressed in wildtype B73 plants (Supplemental Figure 13a) (Whipple et al. 2010) (Xiao et al. 2022). Similarly, editing of three ARF TGTC motifs within the highly conserved 3’UTR of BIF2, the maize homolog of Arabidopsis *PINOID*, resulted in ears with defective axillary meristem initiation reminiscent of previously described *bif2* coding region mutants (Supplemental Figure 13b) (McSteen et al. 2007).

We next investigated whether *cis*-regulatory variation at distal regulatory regions also contributed to phenotypic differences. To this end, we performed CRISPR-based genome editing on the long-range DICE enhancer, known to contribute to herbivore resistance (Zheng et al. 2015). The DICE enhancer lies 143kb upstream of the *BX1* gene, which encodes the first enzyme in the production of the herbivore resistance compound DIMBOA (Frey et al. 1997). Previous data has shown that Mo17, which has higher levels of *BX1* mRNA and higher levels of DIMBOA relative to B73, contains partially duplicated sequence within the genetically mapped DICE enhancer (Figure 5b) (Zheng et al. 2015). DAP-seq data near this region revealed the presence of two conserved DAP-CRMs in B73 and Mo17 (pink boxes in Figure 5a), and an additional DAP-CRM in Mo17. The additional Mo17 DAP-CRM (CRM119799) contained twelve TFs and overlapped with a 3.4kb Mo17-specific insertion (purple region in Figure 5a) (Zheng et al. 2015). Interestingly, our TF binding site data revealed that the Mo17-specific DAP-CRM contained a similar configuration of TFs to that seen in syntenic B73-Mo17-CRM119798, suggesting the increased *BX1* expression seen in Mo17 was caused by a tandem enhancer duplication consisting of at least nine TF binding sites (Figure 5a, Supplemental Figure 14a).

**Figure 5.**
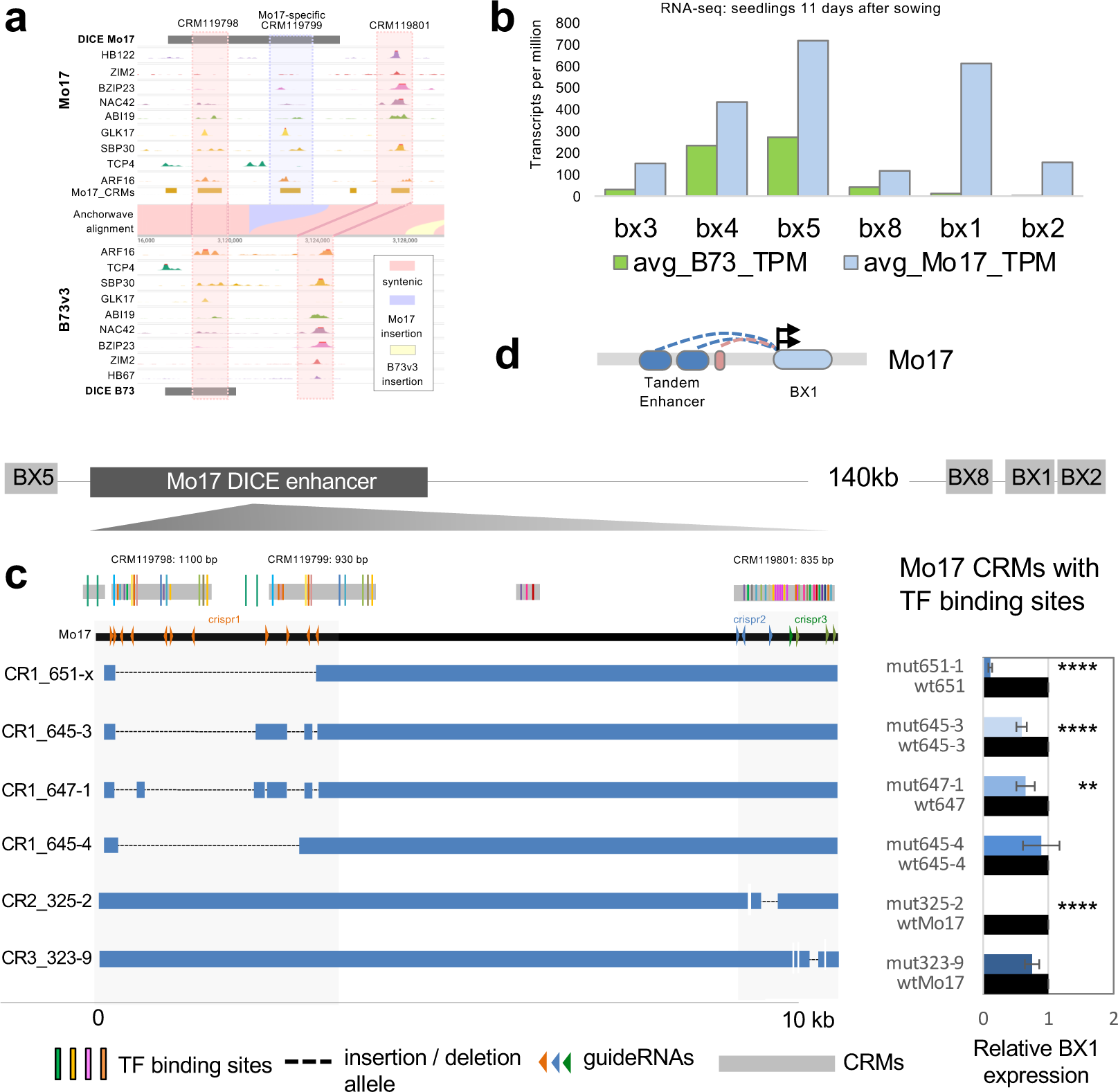
CRISPR induced *cis*-regulatory variation drives expression and phenotypic differences. **a**. JBrowse 2 genome browser screenshot of ∼12kb region surrounding the DICE enhancer in Mo17 and B73v3. Comparative DAP-seq data near the DICE enhancer reveals two conserved DAP-CRMs (pink highlighted area) and one Mo17-specific CRM (purple highlighted area) that appears to be a partial segmental duplication of the upstream CRM and binding sites (CRM119798). **b**. RNA-seq data from 11-day old seedlings from Zhou et al., 2019 showing expression levels (TPM: transcripts per million) of various *BX* genes located near the DICE enhancer. Gene order is same as on chromosome. Mo17 shows 51-fold greater levels of *BX1* expression relative to B73. **c**. CRISPR editing of Mo17 sequences using multiplexed guides near the DICE enhancer revealed specific TF binding sites important for *BX1* expression. Relative *BX1* qRT-PCR expression for six independent alleles is shown on right. Error bars represent standard deviation. **** adjusted pvalue <0.0001, ** adjusted pvalue <0.001. **d**. Schematic depicting individual enhancer components that contribute to enhanced expression of *BX1* in Mo17.

We used three independent CRISPR constructs to generate an allelic deletion series targeting various DAP-seq peaks within the CRMs in Mo17 (Figure 5c). *BX1* expression in the various alleles was measured relative to Mo17 or wildtype sibling controls using qRT-PCR in leaves of 9 to 11-day old seedlings. A large 4.1kb deletion that removed 16 TF binding sites within CRM119798 and CRM119799 caused a 9-fold reduction in *BX1* expression (allele 651-1), while alleles retaining several of these TF binding sites had only a minor 1.5-1.7-fold reduction in expression levels (Figure 5c). The presence of ARF16, GLK17, and BZIP91 sites may be particularly important as their deletion from both CRMs in allele 651-1 caused the greatest reduction in *BX1* expression relative to other alleles. Small deletions in CRM119801 resulted in altered *BX1* expression, with at least one allele (325-2) showing a substantial reduction in *BX1* levels relative to WT Mo17, suggesting this CRM may also impact the expression of *BX1*. Taken together, these results demonstrate that the underlying cause of the DICE enhancer in Mo17 is likely due to a distal tandem enhancer containing at least nine unique-family TF binding sites that boost *BX1* gene expression (Figure 5d). An additional CRM downstream of DICE, may also substantially contribute to *BX1* expression.

Overall, our data show that i) our DAP-seq approach identified biologically relevant regulatory regions; ii) functional TF binding sites can be located within promoters, 3’ UTRs, or distal enhancers; and iii) that knowledge about the exact location of these sites can be used to modulate agricultural traits.

## Discussion

Numerous strategies have been used for mapping non-coding regulatory regions in plants, each with different advantages (Marand et al. 2023) (Marand et al. 2021)(Hajheidari and Huang 2022) (Parvathaneni et al. 2020) (Oka et al. 2017) (Savadel et al. 2021) (Engelhorn et al. 2023; Cahn et al. 2024; Sun et al. 2020). Here, we present a TF-centric approach to annotate the maize non-coding space, taking advantage of several aspects of DAP-seq including the power to profile large numbers of TF family members in multiple genomes, and the ability to assess TF-DNA binding in the absence (or presence) of partner TFs without interference from other *in vivo* factors. Integrating this information with existing non-coding space annotations allowed us to identify individual and composite TF binding sites within both proximal and distal regions that overlapped with GWAS hits and could be used for further functional analysis and targeted breeding. In addition, using DAP-seq we observed direct TF binding by a plant-specific sub-clade of the BHLH family previously not known to bind DNA, while doubleDAP-seq allowed us to tease apart differences in DNA binding specificities for different homo- and heterodimeric combinations of BHLH sub-clade members. Given the large number of TFs known to heterodimerize in protein-protein interaction assays, our results imply that many more novel DNA binding preferences are yet to be revealed.

Our comparative analysis of TF binding in B73 and Mo17 revealed a strong degree (∼64% average) of TF binding site conservation among diverse inbred lines, highlighting the reproducible nature of our data and implying evolutionary constraint likely related to key regulatory functions. Indeed, our data also showed conservation of individual TF binding-target gene pairs between maize and Arabidopsis (Figure 1b), as has been observed for other distantly related plant species (Hendelman et al., 2021). At the same time however, we also observed widespread differences in TF binding across B73 and Mo17, which suggests certain TF binding sites (such as the MADS69 binding site within the *vgt1* QTL) could be important contributors to trait diversity. While structural variants were the largest contributor to genotype-specific differences, SNPs, indels and positional variants were also prevalent and associated with changes in gene expression. Pan-genome analysis of maize genes has revealed that a core set of genes are conserved in related accessions, where on average 67% of genes from a single inbred belong to the core or near-core pan-genome set; (Hufford et al. 2021). These data are similar to the 64% of conserved TF binding sites we observed across B73 and Mo17 although this value would be expected to increase as a greater number of inbred lines are analyzed. Expanding our TF binding site catalog to include sites in additional phenotypically diverse maize inbred lines would help gain a better understanding of the core cistrome and likely help to broadly resolve additional complex QTL and GWAS hits. Our comparative TF binding data involving just two inbreds facilitated molecular clarification of two genetically defined QTL, *vgt1* and DICE, opening possibilities for engineering of desirable agronomic traits such as flowering time and herbivore resistance. Large scale *cis*-regulatory mapping studies will enable a better understanding of how TF binding sites influence gene expression and ultimately phenotype, facilitating the breeding of agronomically robust varieties.

## Methods

### Genomic DNA library constructions, DAP-seq, and DoubleDAP-seq

Genomic DNA libraries were constructed using DNA purified from the aerial portion of 14-day maize seedlings as described in Bartlett et al., 2016). Briefly, 5ug of purified DNA was sheared to 200bp fragments using a Covaris S2. Sheared DNA was bead cleaned with Ampure XP beads (Beckman-Coulter) at a 2:1 bead to DNA ratio. Samples were then end-repaired using the End-It kit (Lucigen), A-tailed with Klenow 3’-5’ exo-(NEB), and truncated Illumina adapters were added with T4 DNA ligase (NEB) overnight at 16 degrees C. Adapter-ligated libraries were bead cleaned with Ampure XP beads at a 1:1 bead to DNA ratio to remove unincorporated adapters and quantified with a Qubit HS kit (Thermo-Fisher).

TF clones were largely obtained from the maize TF clone collection (Burdo et al. 2014). Similarly, all TF family names are based on the Grassius nomenclature (Burdo et al. 2014) (Yilmaz et al. 2009). Please refer to Supplemental Table 1 for corresponding gene IDs. DAP-seq experiments were carried out as described previously (Bartlett et al., 2016) (Galli et al. 2018). Briefly, Gateway pENTR clones from the maize TF collection were LR recombined into the pIX-HALO::ccdB vector (Bartlett et al., 2017) and 1ug of plasmid DNA was used for *in vitro* protein expression using the TNT rabbit reticulocyte expression system (Promega) per the manufacturer’s instructions. *In vitro* protein reactions were incubated for 2 hours at 30 degrees C. HALO-TF protein was subsequently incubated with 10 ug of MagneHALO beads (Promega) for 1 hour rotating at room temperature. Beads were subsequently washed with 100 ul of wash buffer (PBS and 0.005% NP40) three times prior to addition of 1 ug of maize adapter-ligated genomic library diluted in wash buffer. Samples were then rotated for 1 hour at room temperature. Unbound DNA was washed away with six to eight washes of 100 ul of wash buffer, and bound DNA was eluted in 30 ul of Elution Buffer (EB, 10mM Tris). Sample were heated to 98 degrees C for 10min, placed on ice, and the eluted DNA was recovered from the beads and stored at -20 degrees C prior to performing PCR enrichment and barcoding. Eluted DNA was PCR amplified as described in Bartlett et al., 2016 using 19 cycles and dual indexed Illumina TruSeq primers. Samples were sequenced with either the NextSeq550 (Single end, 75bp reads), NovaSeq6000 (paired end, 150bp reads), or HiSeqX (paired end, 150bp reads).

DoubleDAP-seq experiments were carried out by simultaneously expressing pIX-HALO-TFs and pIX-SBPtag clones (Li et al. 2023) in a 50 ul TNT rabbit reticulocyte reaction containing 1 ug of each plasmid in a 2 hour incubation at 30 degrees C. Subsequently 1 ug of adapter-ligated library was diluted in wash buffer and added to the reticulocyte reaction together with 10 ul of MagneHALO beads for a final volume 100 ul. Samples were rotated for 2 hours at room temperature, washed eight to ten times in wash buffer with the use of a magnet. Bound DNA was eluted with 30 ul of EB and incubation at 98 degrees C for 10min. Sample enrichment and sequencing was performed as for single DAP experiments.

### Read mapping and blacklist construction

Reads were quality trimmed using trimmomatic (Bolger, Lohse, and Usadel 2014) and mapped to either the B73v5 or Mo17 CAU1.0 genomes with bowtie2 using default parameters (Langmead and Salzberg 2012). Mapped reads were filtered to retain only reads with MAPQ greater than 30 using ‘samtools view -q 30’. Stringent criteria were established to exclude artifactual binding regions by the generation of a blacklist that captured the majority of non-specific peaks. A list of sites bound in nearly all TF datasets and the negative control HALO-GST sample (Galli et al. 2018) was manually curated for both B73v5 and Mo17 CAU1.0 and can be found in Supplemental Tables 3 and 4.

### Peak calling analysis and target gene assignment

Peaks were called with GEM3 (Guo, Mahony, and Gifford 2012) using a standard threshold method of adjusted p-value of 0.00001 (--q 5). For datasets that produced two few peaks at this threshold (i.e. <10,000), the default threshold of 0.01 (--q 2) was used. For datasets that exceeded 100,000 peaks at the q5 threshold, a q10 threshold (e-10) was applied. The total coverage of peaks genome-wide was calculated using a 30bp region surrounding the peak summit using the bedtools slop and merge utilities (Quinlan and Hall 2010), and dividing by the total of size of B73v5 chromosomes 1-10. Putative target genes were assigned to each peak using ChIPseeker and default assignment priorities (Yu, Wang, and He 2015). Promoters were defined as -10,000bp to +1bp relative to the TSS where gene features were annotated according to the Zm-B73-REFERENCE-NAM-5.0_Zm00001eb.1.gff3.gz annotation file downloaded from maizeGDB.

### Motif analysis

The GEM events reported were ranked by q-value then by fold enrichment. Sequences for 201 bp region centered at the top 1000 GEM events were extracted from the corresponding genome (B73v5 or Mo17 CAU1.0) and used in *de novo* motif discovery by meme-chip (version 5.3.0) with the parameter “-meme-searchsize 0” (Machanick and Bailey 2011). For comparing motif scores between the B73-specific peaks and the syntenic Mo17 regions, aligned sequences between the two genomes were extracted from the Anchorwave .maf output file using the UCSC utility mafInRegions and scored against the PWM motifs from meme-chip using MAGGIE (Shen et al. 2020).

### Heterodimer model analysis for DoubleDAP-seq

Protein heterodimers of BHLH125 and BHLH85 were modeled using AlphaFold on Google CoLAb Pro (Jumper et al. 2021) (Mirdita et al. 2022). Protein input sequences were the same as those used for DAP-seq clones and did not match those present in the AlphaFold database which had the incorrect sequence for ZmBHLH85.

### GWAS analysis

GWAS summary statistics computed by the NAM genomes project were downloaded from CyVerse Data Commons: /iplant/home/shared/NAM/NAM_genome_and_annotation_Jan2021_release/SUP PLEMENTAL_DATA/NAM-GWAS-PVE-files/SNPs. SNP data was downloaded from /iplant/home/shared/NAM/NAM_genome_and_annotation_Jan2021_release/SUP PLEMENTAL_DATA/NAM-SV-projected-V8. Linkage disequilibrium (LD) was computed for only biallelic SNPs using PLINK (version 1.90b6.21) with the parameters ‘--make-founders --r2 dprime --ld-window-kb 100 --ld-window 100000 --ld-window-r2 0.2’. The GWAS functional enrichment tool GARFIELD (Iotchkova et al. 2019) was used to annotate LD-pruned SNPs (LD r2 > 0.1) by their overlap with DAP-seq peaks of each TF. Subsequently, the odds ratio and significance of the overlap were computed for various GWAS significance levels, accounting for minor allele frequency, distance to the nearest TSS, and the number of LD proxies (r2 > 0.8).

### Whole genome alignment analysis and coordinate liftover

A whole genome alignment of the B73v5 and Mo17 genomes were performed using Anchorwave as described in Song et al., 2021. The .maf output file was then converted to a .chain file using the maf-convert script of LAST 1296 (Kielbasa et al. 2011). Coordinate liftover was performed using Crossmap (Zhao et al. 2014) and the chain files generated from Anchorwave and maf-convert of LAST. Visualization was performed using IGV (single genome view) (Robinson et al. 2011) and JBrowse 2 (Diesh et al. 2023) (comparative view using the chain files generated by the maf-convert script (LAST) from the Anchorwave .maf file).

### Normalization of TF diversity panel for determining genotype-specific peaks

BAM files of mapped reads were downsampled using samtools -s to adjust the number of mapped reads between Mo17 and B73 to the lower value of the two datasets for the same TF. Peak calling was then performed on the downsampled datasets using GEM3 (Guo, Mahony, and Gifford 2012) and the threshold was adjusted to achieve similar number of peaks between Mo17 and B73. Peak coordinates were then ‘lifted’ using CrossMap (Zhao et al. 2014). Pearson correlation between coordinate-lifted and reference peaks was determined using the deeptools2 multiBigWigsummary utility (Ramirez et al. 2014). Overlap between CrossMap lifted peak coordinates and the reference genotype was determined using bedtools2 (Quinlan and Hall 2010). Shared and genotype-specific peaks were determined for each TF by taking the top 20% of peaks from one genotype and comparing them to the total peaks of the opposing genotype to reduce false positives and negatives resulting from thresholding differences arising during peak calling. Positional variation (posV) relative to the TSS was calculated as follows: B73v5 gene models in gff3 format were converted to Mo17 genome coordinates using LiftOff (Shumate and Salzberg 2021) with default parameters. Putative target genes were assigned using ChIPseeker (Yu, Wang, and He 2015) using either the B73v5 gene models (B73 datasets) or B73v5 gene models lifted to Mo17 (Mo17 datasets). This ensured differences in distance to TSS were not due to differences in gene model annotation. Differences in distance to TSS were then calculated for those shared peaks with the same assigned putative target gene. Only positional variants for B73 peaks were determined but similar percentages would be expected for Mo17 peaks.

To assess peak overlap with SNPs, indels (less than 50bp), and structural variants (here defined as indels greater than 50bp), a genome-wide VCF file was generated using SyRI (Goel et al. 2019) as follows. First, the .maf output file from Anchorwave (B73v5 as reference, Mo17 as query) was converted to .sam format using the maf-convert script of LAST (Kielbasa et al. 2011). This file was then used with SyRI to generate the B73-Mo17 .VCF file based on the following parameters: syri -c anchorwave_Mo17toB73v5.sam -F S --prefix anchorwave_Mo17toB73v5_sam_ --cigar -f --log DEBUG -r Zm-B73-REFERENCE-NAM-5.0.id_chrs_mg.fa -q Mo17_CAU-1/Zm-Mo17-REFERENCE-CAU-1.0.id_chr_nuc_mg.fa. The resulting VCF file was parsed to separate SNPs, small indels less than 50bp, structural variants (indels greater than 50bp), ‘not aligned’ sequence, and duplicated regions. The bedtools intersect utility from the BEDtools2 suite (Quinlan and Hall 2010) was used to quantify overlap with variant features. Peaks and features were considered overlapping if their coordinates shared greater than or equal to 1bp.

### Differential Gene expression correlation and root cell-type specific analysis

Raw RNA-seq reads for the selected tissues in B73 and Mo17 were downloaded from NCBI SRA (PRJNA482146; Supplemental Table 5. Transcript quantification was done using the RSEM software package (version 1.3.3) (Li and Dewey 2011) with the STAR aligner (version 2.7.6a) (Dobin et al. 2013). RSEM references were built using the Mo17 and B73v5 genome sequences with annotations of genes mappable between Mo17 and B73. To create the Mo17 gene annotation, the B73v5 gene model (Zm-B73-REFERENCE-NAM-5.0_Zm00001eb.1.gff3) was mapped from the reference genome sequence Zm-B73-REFERENCE-NAM-5.0 to the target genome sequence Zm-Mo17-REFERENCE-CAU-1.0 using liftoff (version v1.6.3) (Shumate and Salzberg 2021). The resulting annotation file was used with the Mo17 genome sequence to build the Mo17 RSEM reference. To create the B73 gene annotation, genes in the B73v5 gene model were filtered to keep only genes that were lifted to Mo17. The resulting annotation file was used with the B73v5 genome sequence to build the B73 RSEM reference. Gene expression values were then calculated from the paired-end RNA-seq read files using the corresponding RSEM reference genomes. The RSEM results were imported into R by tximport (version 1.30.0) (Soneson, Love, and Robinson 2015) for differential expression analysis by DESeq2 (version 1.42.1). To remove genes with low counts in a majority of the samples, pre-filtering was performed by keeping genes that had read counts of at least 10 in a minimum number of samples, where the minimum was set to the lowest number of replicates for the two genotypes in each tissue (Supplemental Table 5). For each tissue, differential gene expression between B73 and Mo17 was computed using the standard DESeq analysis settings. DEGs were identified as genes that had absolute log2 fold change greater than 2 and FDR adjusted P-value lower than 0.05.

Putative shared or B73-specific target genes were assigned by ChIPseeker (Yu, Wang, and He 2015) based on genes that have shared or B73-specific peaks within -10kb to +300 bp from the TSS in B73 gene model Zm-B73-REFERENCE-NAM-5.0_Zm00001eb.1.gff3. Fisher’s exact test was used to evaluate the significance of overlap between B73 vs. Mo17 DEGs and genes associated with shared and B73-specific DAP-seq peaks. Two-by-two contingency tables (DEG vs. DAP-seq target) were created for the shared and B73-specific target genes of the TFs in the diversity panel and for all nine tissue types,which were used in one-sided Fisher’s exact tests by the fisher.test function in R. The reported P values were then adjusted for multiple hypothesis testing by the Benjamini–Hochberg procedure and transformed to -log10 scale for plotting by the R package ComplexHeatmap (Gu, Eils, and Schlesner 2016) (Gu 2022). Row and column clustering were done using Euclidean distance and complete linkage methods.

For overlap with root cell type markers, the list of cell type-specific marker genes for maize weas obtained from Supplementary Table 4 of Guillotin et al. (2023). The root cell-type specific markers for 19 clusters from single cell RNA-seq were converted from B73v4 gene model to v5 using the liftoff files provided by MaizeGDB (https://ars-usda.app.box.com/v/maizegdb-public/folder/165362280830). Fisher’s exact test was performed to test the overlap between the cell type-specific marker genes and the shared and B73-specific target genes. The reported P values were adjusted for multiple hypothesis testing by the Benjamini–Hochberg procedure and transformed to -log10 scale for plotting by the R package ComplexHeatmap (Gu, Eils, and Schlesner 2016) (Gu 2022).

To determine whether positional variation (posV) among shared promoter peaks of B73 and Mo17 was associated with differential gene expression, the posV peaks for 27 TFs were analyzed, wherein the DAP-seq TF targets were divided in two groups: posVs greater than 500bp and posVs less than 500bp. Considering only the DAP-seq TF targets that were positionally variant between inbreds, two-by-two continency tables (a gene is DEG or not DEG vs. a target gene is associated with posV greater than 500bp or less than 500bp) were created for each of the 27 TFs and for all nine tissue types, which were used in one-sided Fisher’s exact tests by the fisher.test function in R. The reported P values were then adjusted for multiple hypothesis testing by the Benjamini–Hochberg procedure.

### Orthogonal dataset overlap analysis

The MOA-seq data from ear tissue (Savadel et al. 2021) remapped to B73v5 was obtained from maizeGDB. ATAC-seq data from leaf and ear were obtained from Ricci et al., 2019 and Hufford et al., 2021 via maizeGDB. Histone modification data was from Ricci et al. 2019. NAM consortium UMR data was obtained from maizeGDB (Hufford et al., 2021). Bed annotation files of B73v5 transposable elements were obtained from maizeGDB. The conserved non-coding sequence (CNS) regions from sorghum were obtained from Song et al. 2021 with B73v4 coordinates converted to B73v5 coordinates with the ensemblplants Assembly Converter tool. Overlap analysis was performed using the bedtools intersect tool from the BEDtools2 suite (Quinlan and Hall 2010). Peaks were considered to overlap with a particular feature if their coordinates shared greater than or equal to 1bp. Mo17 UMR and ATAC-seq peak data shown in JBrowse 2 genome browser screenshots was from Noshay et al. 2021.

### GO enrichment analysis

GO enrichment was performed using the ShinyGO server (Ge, Jung, and Yao 2020). Analysis was performed by selecting top peaks (according to signal value) that were located in UTRs, putative promoters (1-10kb from TSS), and downstream (<300bp from TES) regions (excluding distal, i.e. >10kb from TSS, exonic and intronic regions) to focus datasets on those gene most likely relevant to our analysis. In most cases, putative target genes associated with the first 3000 peaks were used for GO analysis. In the event when these did not produce significant results, the top 6000 peaks were used.

### TF diversity panel

This panel consisted of 66 TFs and TF combinations (i.e. trimers of NFY-A, NFY-B, and NFY-C, and dimers of BHLH67/BA1 and ZmSPTs discussed above) from this study and several previously published maize DAP-seq datasets (Galli et al. 2018) (Ricci et al. 2019) (Dong et al. 2020) (Dai et al. 2022) (Wu et al. 2023) (Bang et al., 2024). Only datasets that showed unique binding motifs, belonged to a distinct subfamily, and/or had a Pearson correlation less than 0.5 were included.

### CRIPSR constructs

For CRISPR constructs targeting *cis*-regulatory regions near *TSH1* and the DICE enhancer, multiplexed guides were cloned into pBUE411(Xing et al. 2014) using NEB HiFi-based Gibson assembly. Constructs were transformed into maize HiII embryos via Agrobacterium mediated transformation. T0 plants were crossed to B73 (TSH1 and BIF2) or Mo17 (DICE) to generate edits in the B73 and Mo17 backgrounds respectively. Alleles were fixed by removal of the Cas9-guide cassette via genetic segregation and backgrounds were further purified by subsequent backcrossing. For the CRISPR construct targeting the *cis*-regulatory region near the *BIF2* gene, three gRNA cassettes were assembled and cloned them into the pBUE411-GGB vector using the Golden Gate Assembly method (Chen et al., 2021). Subsequently, the resulting construct was transformed into the maize inbred line B104. In the T0 generation, plants with edits at the target region were crossed to B104 once and then were selfed to obtain homozygous edits while simultaneously removing the transfer DNA (T-DNA). It should be noted that a genome assembly error was observed in the B73v5 genome near the DICE locus. The JBrowse 2 screenshot shown in in Figure 5a therefore uses DAP-seq data mapped to B73v3.

### qRT-PCR analysis of DICE edited alleles

Plants were grown in a Conviron growth chamber with 16 hours of light and 8 hours of dark at 26 degrees C. Total RNA was extracted from plants homozygous for each of the CRISPR DICE alleles from the middle of the third leaf on seedlings 9-11 days after sowing. Three biological replicates, each containing three homozygous mutant samples, were harvested for each allele along with three biological replicates of either segregating homozygous wildtype siblings or wildtype parental genotype. cDNA was made using qScript cDNA synthesis kit (QuantaBio). qPCR was performed using PerfeCTa SYBR Green FastMix (QuantaBio) with primers and conditions shown in Supplemental Table 6, using an Eco Real-Time PCR system (Illumina). Analysis was performed using the ddCT method with the qRAT tool and limma statistical framework (Flatschacher, Speckbacher, and Zeilinger 2022; Ritchie et al. 2015).

## Acknowledgements

This work was supported by the NIH award R35GM138143 to S.C.H. and NSF Plant Genome Research Project grant IOS-1916804 to S.C.H. and A.G.

---

**Supplemental Figure 1.**
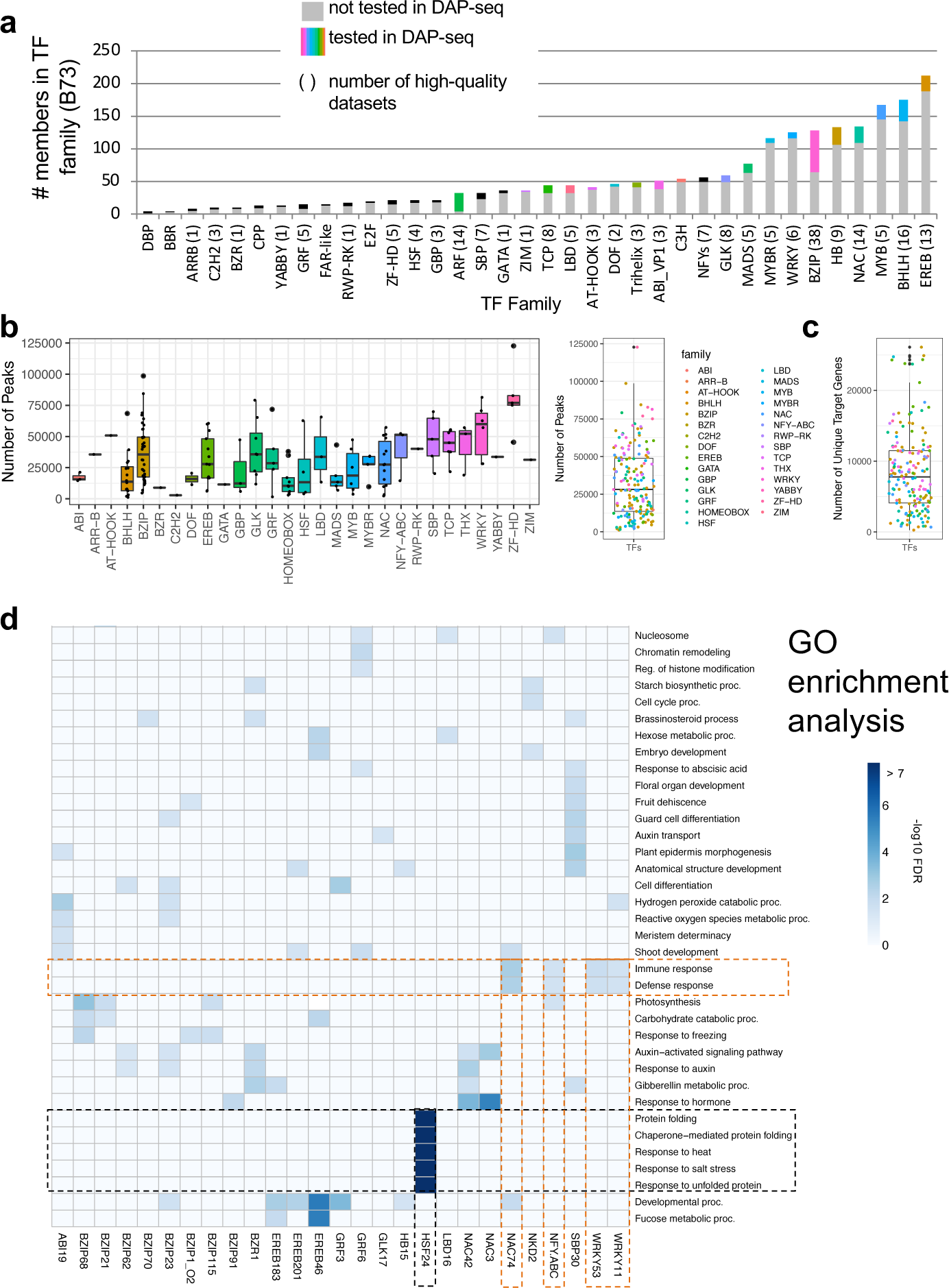
DAP-seq data samples broadly across maize TF families. **a.** Stacked bar graph showing 37 TF families in which at least one family member was tested in DAP-seq. The number of members tested for each is shown with a colored bar; the number of members for which high quality datasets that were obtained is shown in paratheses next to the family name. **b.** Boxplots showing the distribution of the number of peaks obtained for each TF tested in DAP-seq. The left panel shows total number of peaks based on family and the right panel shows total distribution of peaks. Colors indicate different TF families. **c.** Boxplot showing the distribution of the number of target genes obtained for each TF tested in DAP-seq. **d.** GO enrichment analysis of putative target genes assigned to peaks of selected TFs.

**Supplemental Figure 2.**
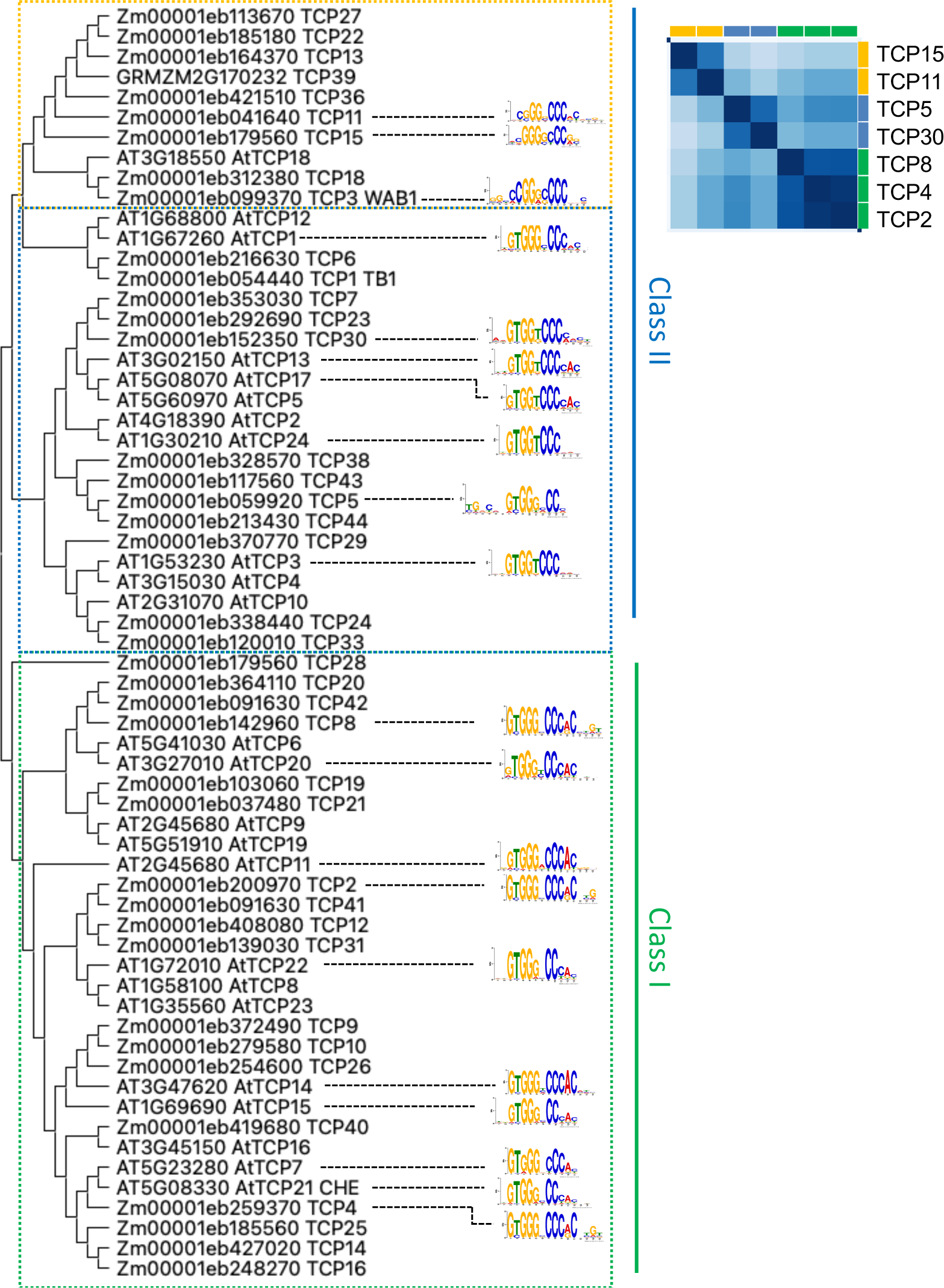
Sub-clade specific DNA-binding motifs and binding preferences of TCP family members. **a.** Maximum likelihood phylogeny of maize and Arabidopsis TCP family members annotated with sub-clade specific binding motifs. Arabidopsis motifs from O’Malley et al., 2016. **b.** Pairwise Pearson correlation of DAP-seq binding profiles between maize TCP family members showing distinct profiles for Class I and Class II TCP family members.

**Supplemental Figure 3.**
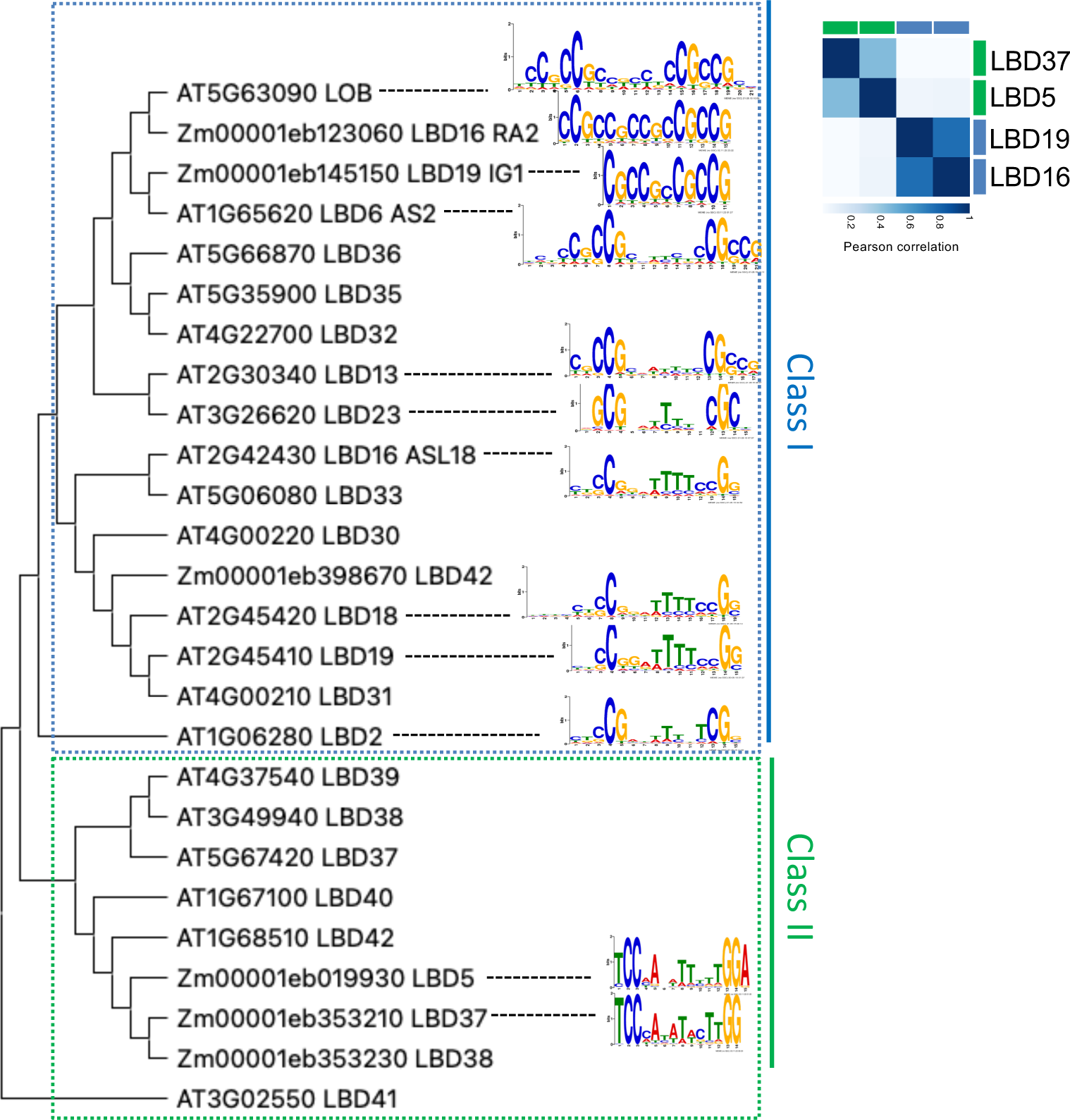
Sub-clade specific DNA-binding motifs and binding preferences of LBD family members. **a.** Maximum likelihood phylogeny of maize and Arabidopsis LBD family members annotated with sub-clade specific binding motifs. Arabidopsis motifs from O’Malley et al., 2016. **b.** Pairwise Pearson correlation of DAP-seq binding profiles between maize LBD family members showing distinct profiles for Class I and Class II family members.

**Supplemental Figure 4.**
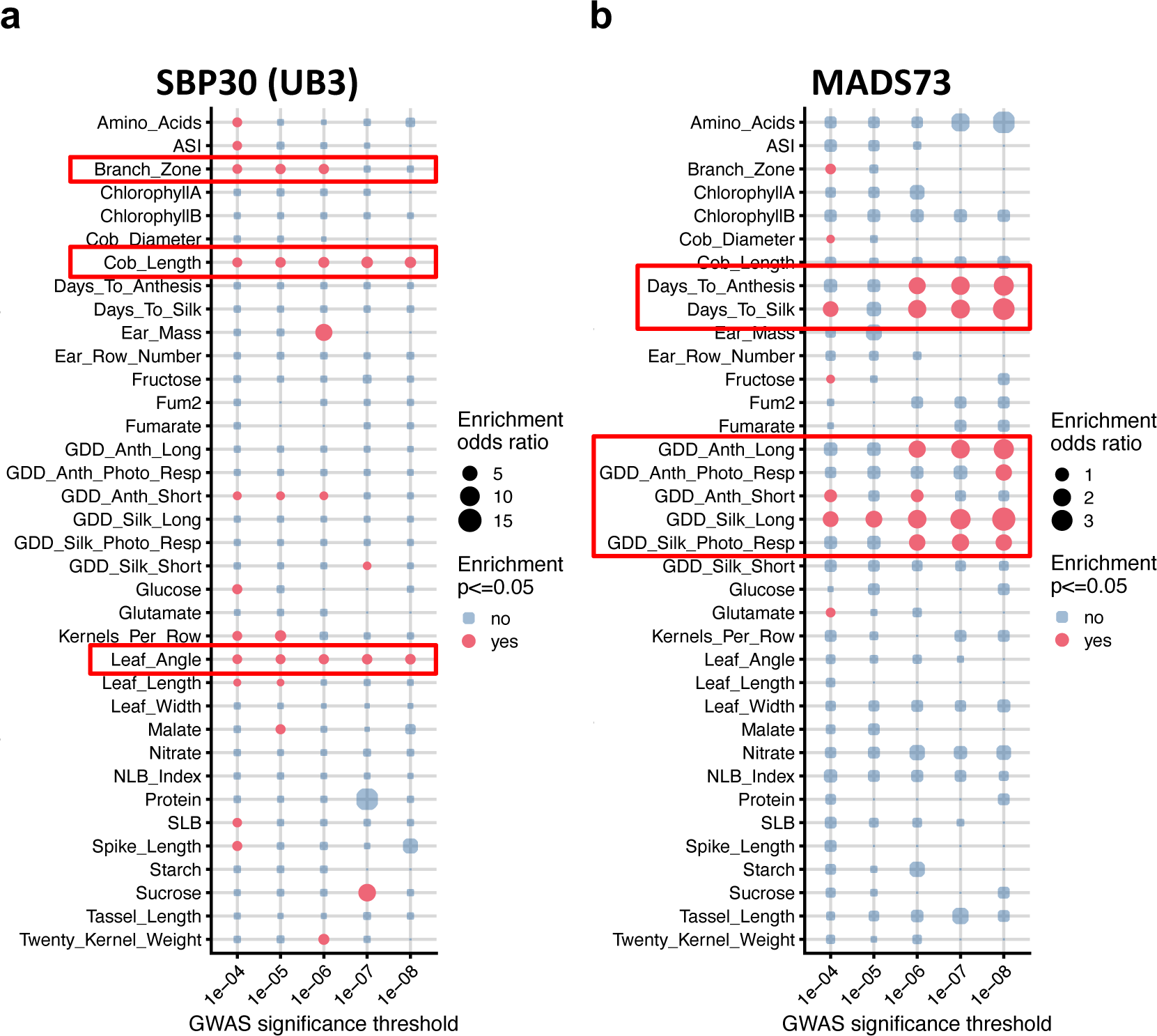
GWAS enrichment analysis of selected TFs and traits from Wallace et al., 2014. **a**. Bubble plots showing enrichment for GWAS SNPs associated with branch zone, cob length, and leaf angle that overlapped SBP30/UB3 DAP-seq peaks. **b**. Bubble plot showing MADS73 DAP-seq peaks are significantly enriched for SNPs associated with several flowering-related traits.

**Supplemental Figure 5.**
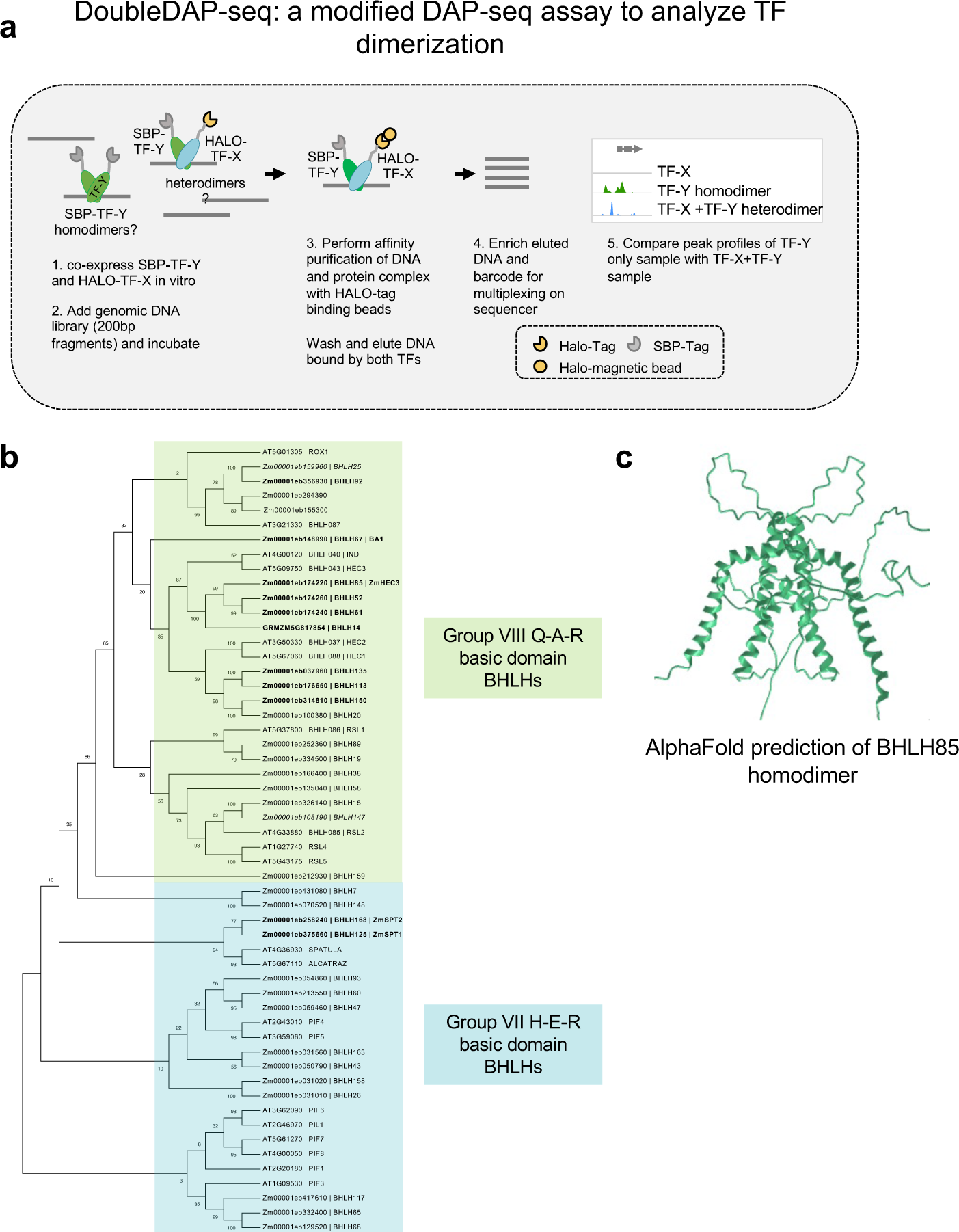
**a**. Schematic showing the doubleDAP-seq assay in which putative heterodimeric TF-DNA complexes can be pulled down and compared to results from single protein DAP-seq assay to assess binding site specificity differences. **b**. Neighbor-joining phylogeny based on amino acid similarity of group VII (blue) and group VIII (green) BHLHs from maize and Arabidopsis. Members that were tested in DAP-seq are show in bold. Three members that were tested in DAP-seq that did not yield any peaks are shown in italics. **c**. AlphaFold prediction of BHLH85 homodimer.

**Supplemental Figure 6.**
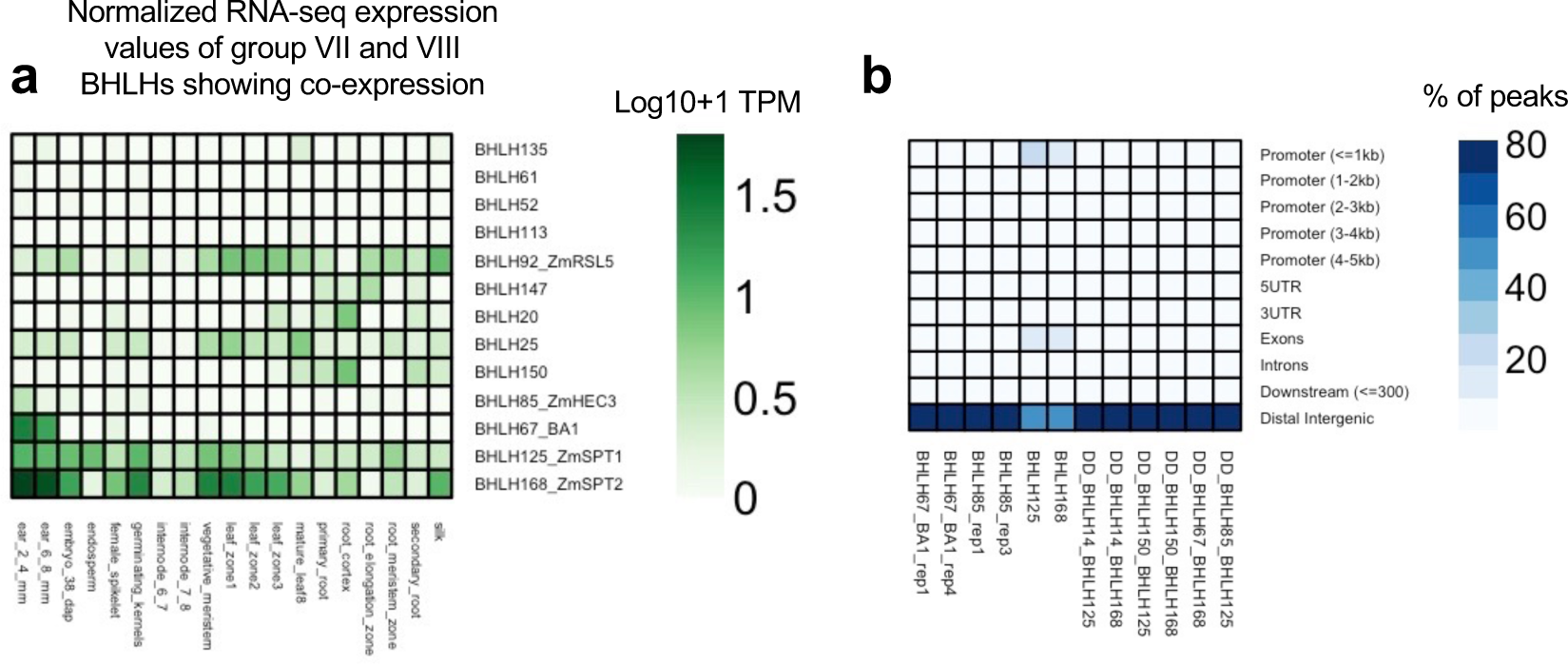
**a.** Normalized RNA-seq expression (TPM; transcripts per million) of group VIII and VII BHLHs showing heterodimer pairs are co-expressed in the same tissues. Normalized expression data from Walley et al., 2016 **b**. Prevalence of binding sites in various gene features differs for group VIII and group VII BHLHs.

**Supplemental Figure 7.**
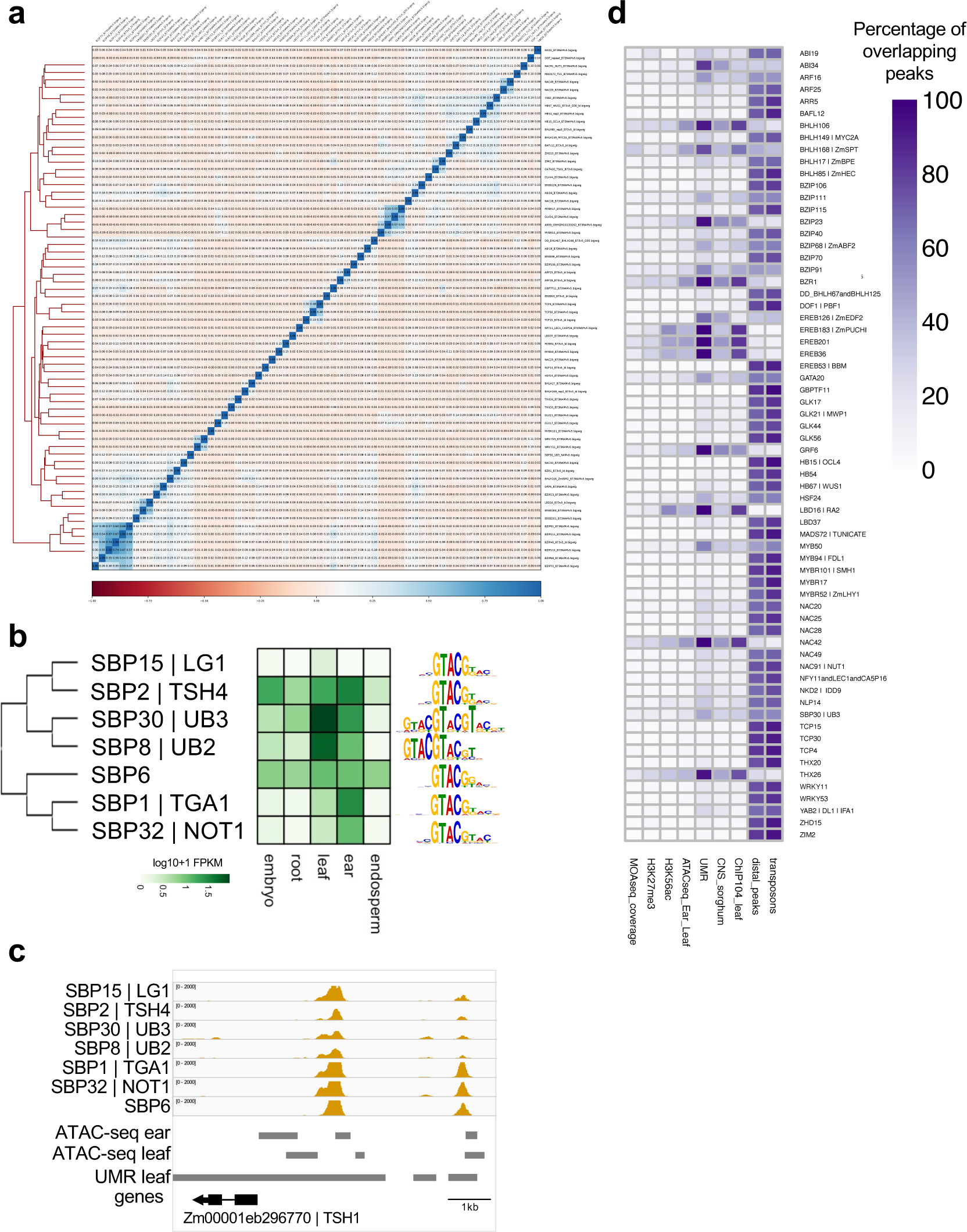
**a.** Heatmap showing Pearson correlation of genome-wide binding events for 66 TFs in the diversity panel. **b**. Phylogeny, tissue-specific RNA-expression (TPM; transcripts per million), and binding motifs of selected SBP TFs tested in maize DAP-seq. **c**. Genome browser screenshot of SBP TFs shown in b at the TSH1 locus. **d**. Heatmap showing the percentage of peaks that overlap various orthogonal functional datasets. See Methods for source data.

**Supplemental Figure 8.**
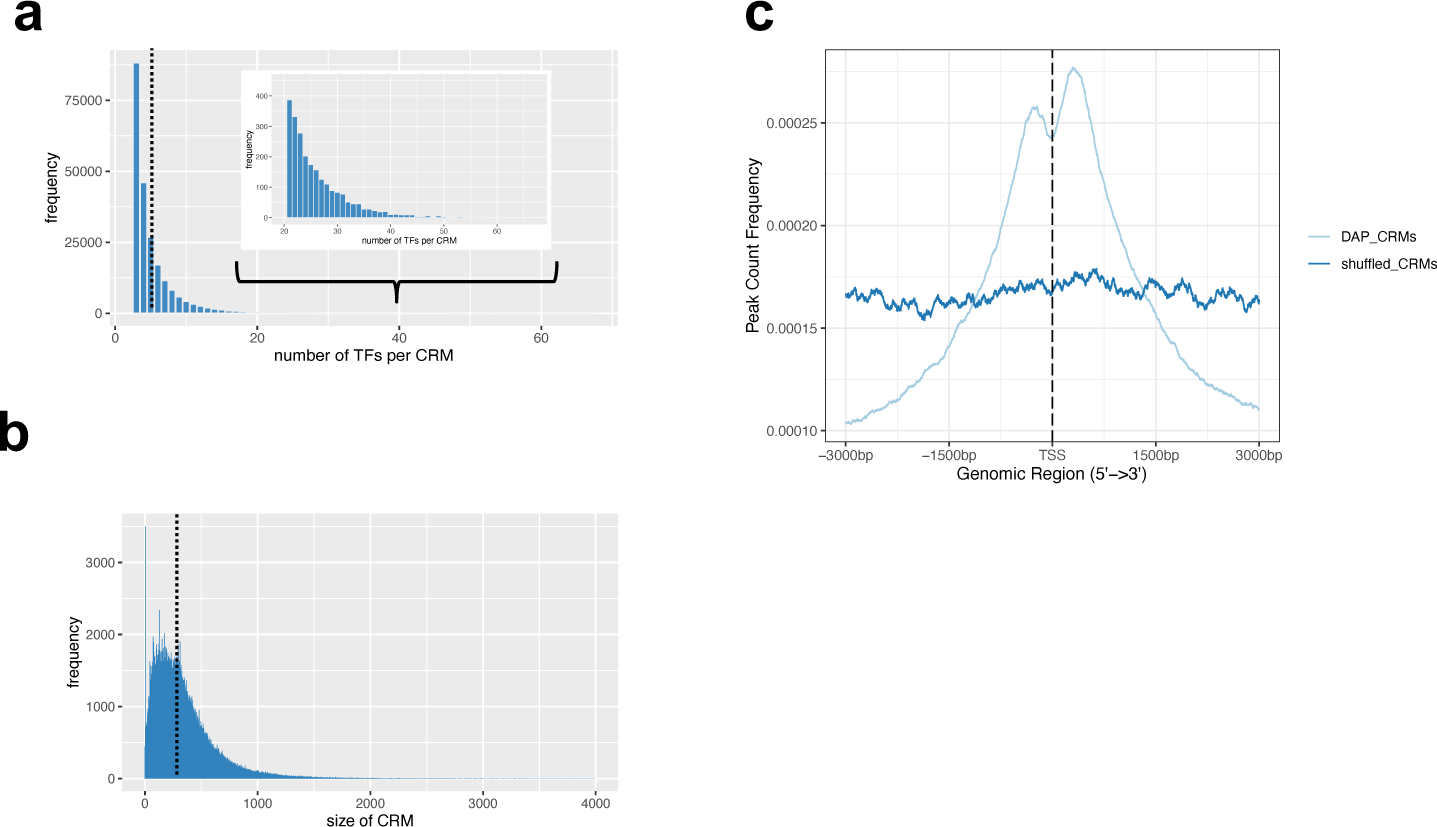
**a.** Histogram of the number of DAP-CRMs containing the specific number of TFs per DAP-CRM. The black dotted line indicates the average number of TFs per DAP-CRM (5.3). The inset shows a close-up on the number of TFs from 20-63 TFs per DAP-CRM. **b.** Histogram showing the size distribution of DAP-CRMs. The dotted line indicates the average size (344bp) of all DAP-CRMs. **c.** Frequency of DAP-CRMs versus randomly shuffled CRMs of the same size within a 3kb window surrounding the TSS.

**Supplemental Figure 9.**
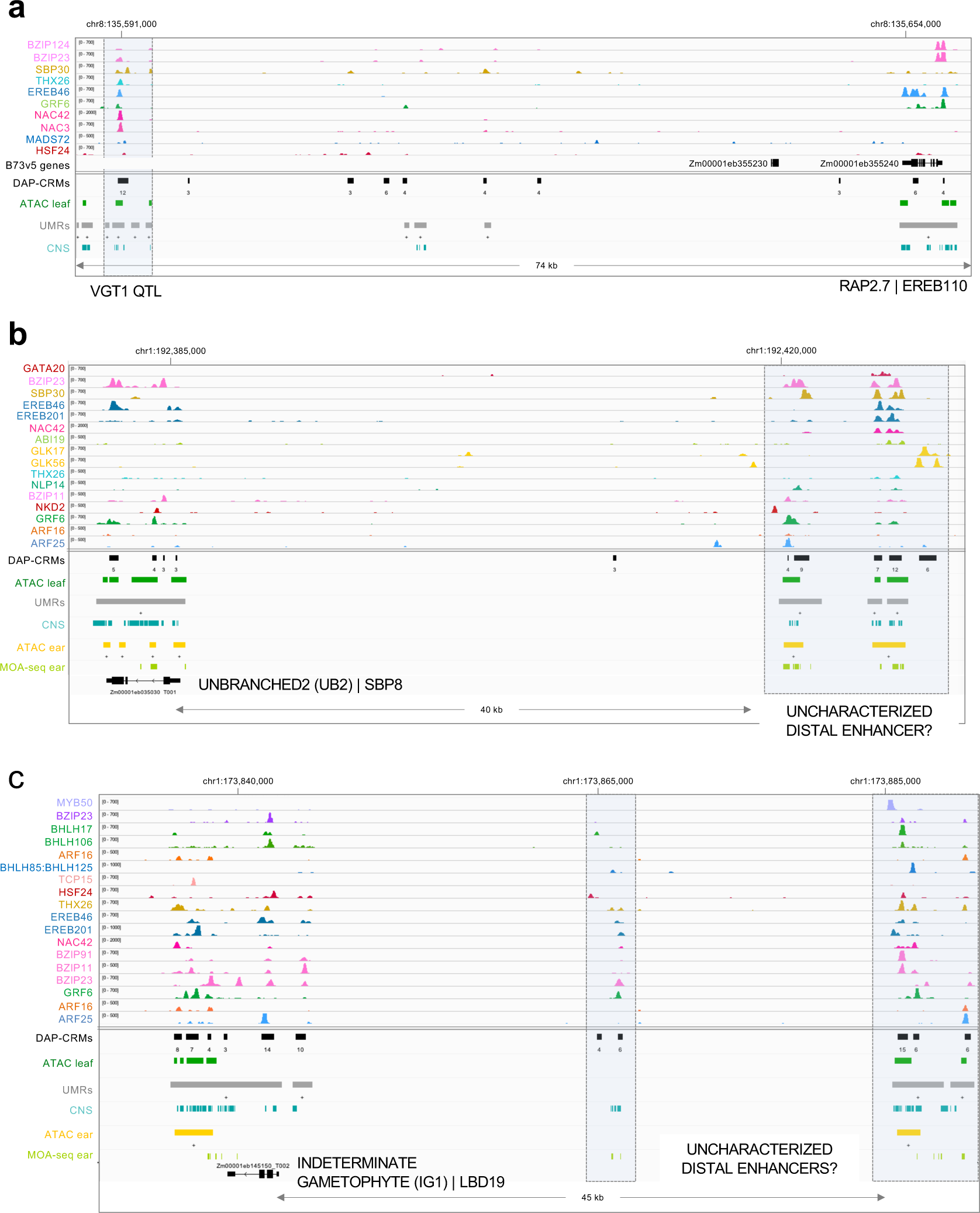
**a**. B73v5 IGV genome browser screenshot of the *vgt1* and *RAP2.7* locus showing binding by DAP-seq TFs. The number below the DAP-CRM track indicates the number of TF peaks present in the DAP-CRM. **b.** Genome browser screenshot of the *TASSEL SHEATH1* (*TSH1)* locus showing binding by DAP-seq TFs. Number below the DAP-CRM track indicate the number of TF peaks present in the DAP-CRM. **c**. Genome browser screenshot of the *INDETERMINATE GAMETOPHYTE1 (IG1)* locus showing binding site location of DAP-seq TFs.

**Supplemental Figure 10.**
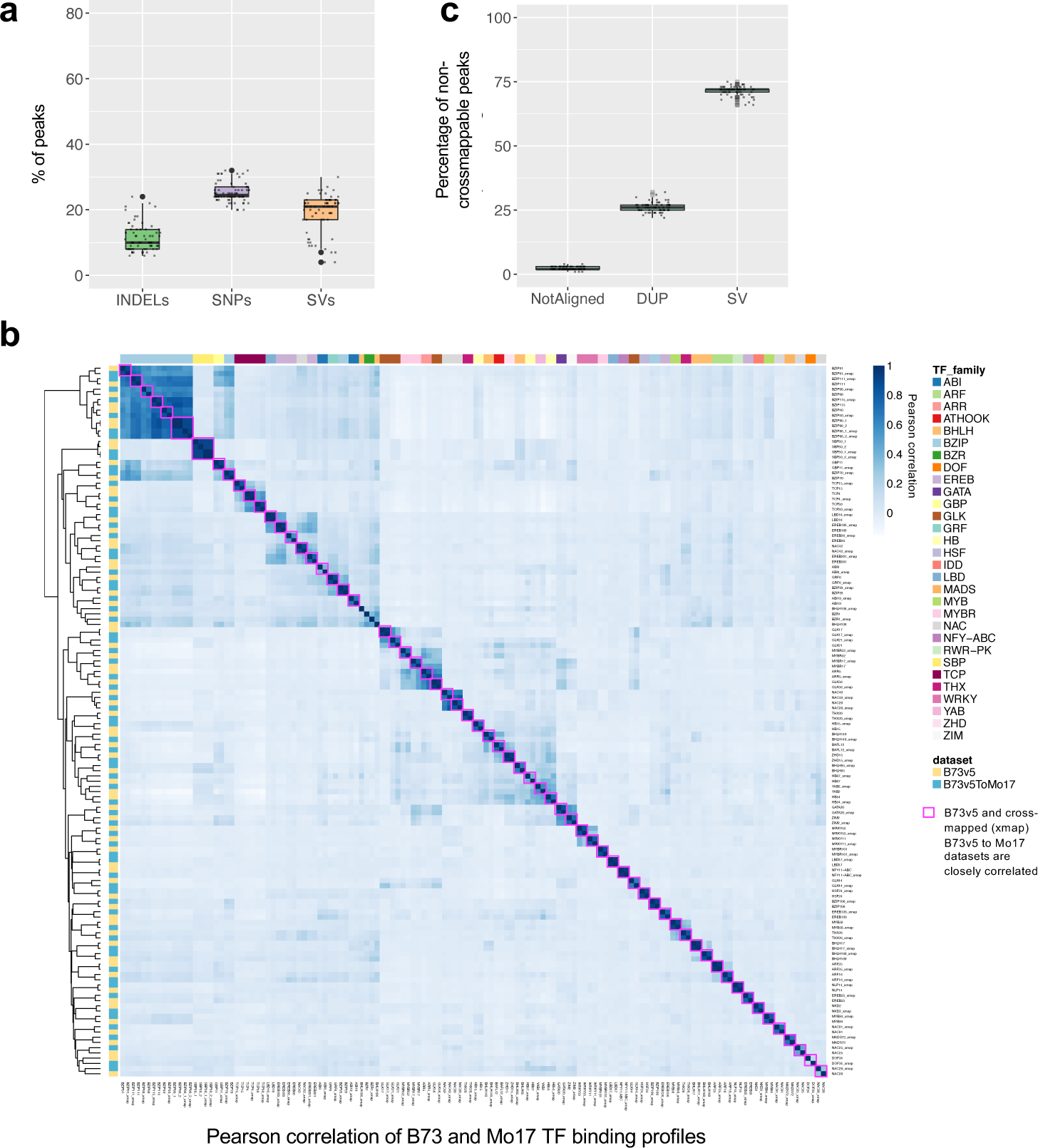
**a**. Percentage of all peaks from the TF diversity panel datasets that overlapped with B73v5-Mo17 SNPs, small indels (less than 50bp), and/or structural variants (indels greater than 50bp). Each datapoint corresponds to the percentage of peaks from an individual TF overlapping the indicated category. **b**. Pearson correlation of B73 and Mo17 TF binding profiles. B73 (side bar in yellow color) and Mo17-lifted peak datasets (side bar in teal color) show high Pearson correlation values. Most datasets cluster most closely with their corresponding dataset from Mo17 (magenta boxes). **c**. Percentage of non-crossmappable peaks (coordinates could not be lifted) that overlapped with ”Non-Alignable” region (NotAligned), duplicated regions where cross-mappability was ambiguous (DUP), or structural variants (SV, insertions or deletions greater than 50bp). Each datapoint corresponds to the percentage of peaks from an individual TF overlapping the indicated category.

**Supplemental Figure 11.**
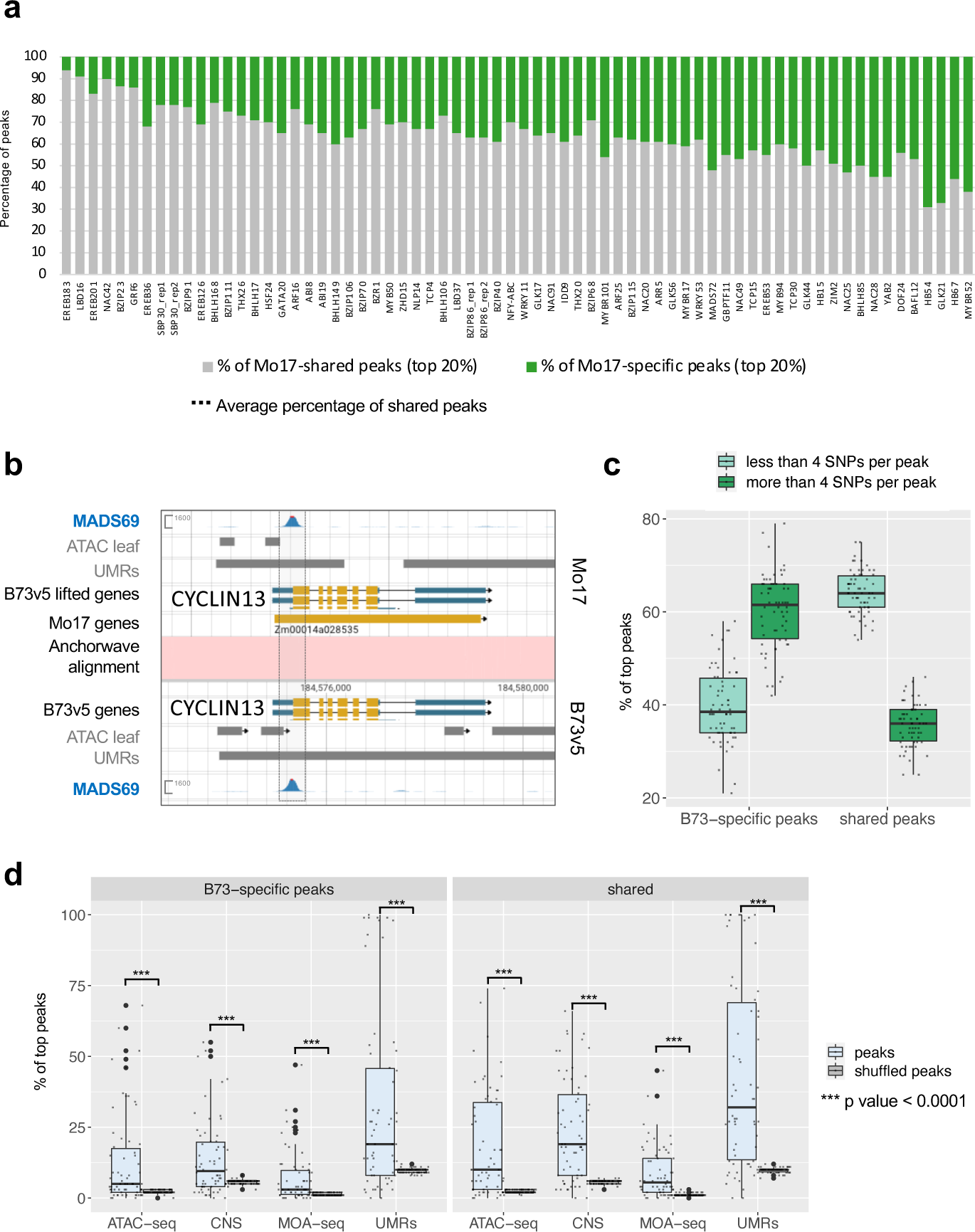
**a**. Percentage of Mo17 TF diversity panel peaks also found in B73 (shared, grey) and those found only in Mo17 but not B73 (Mo17-specific, green). Mo17 datasets show a similar percentage of Mo17-specific and Mo17-B73 shared peaks relative to those seen for B73-specific and B73-Mo17 shared peaks (TF order is same as in Figure 4a). **b**. JBrowse 2 genome browser screenshot showing similar binding intensity of MADS69 in the 5’UTR of *CYCLIN13*. **c**. B73-specific peaks more often contain more than four SNPs per peak. Shared peaks more often contain less than 4 SNPs per peak. **d**. Overlap of B73-specific peaks and B73-Mo17 shared peaks with orthogonal datasets. Both B73-specific peaks and shared peaks show statistically significant enrichment (p value < 0.0001, Student’s t test) relative to randomly shuffled peaks, providing support that both B73-specific peaks and shared peaks could be functional.

**Supplemental Figure 12.**
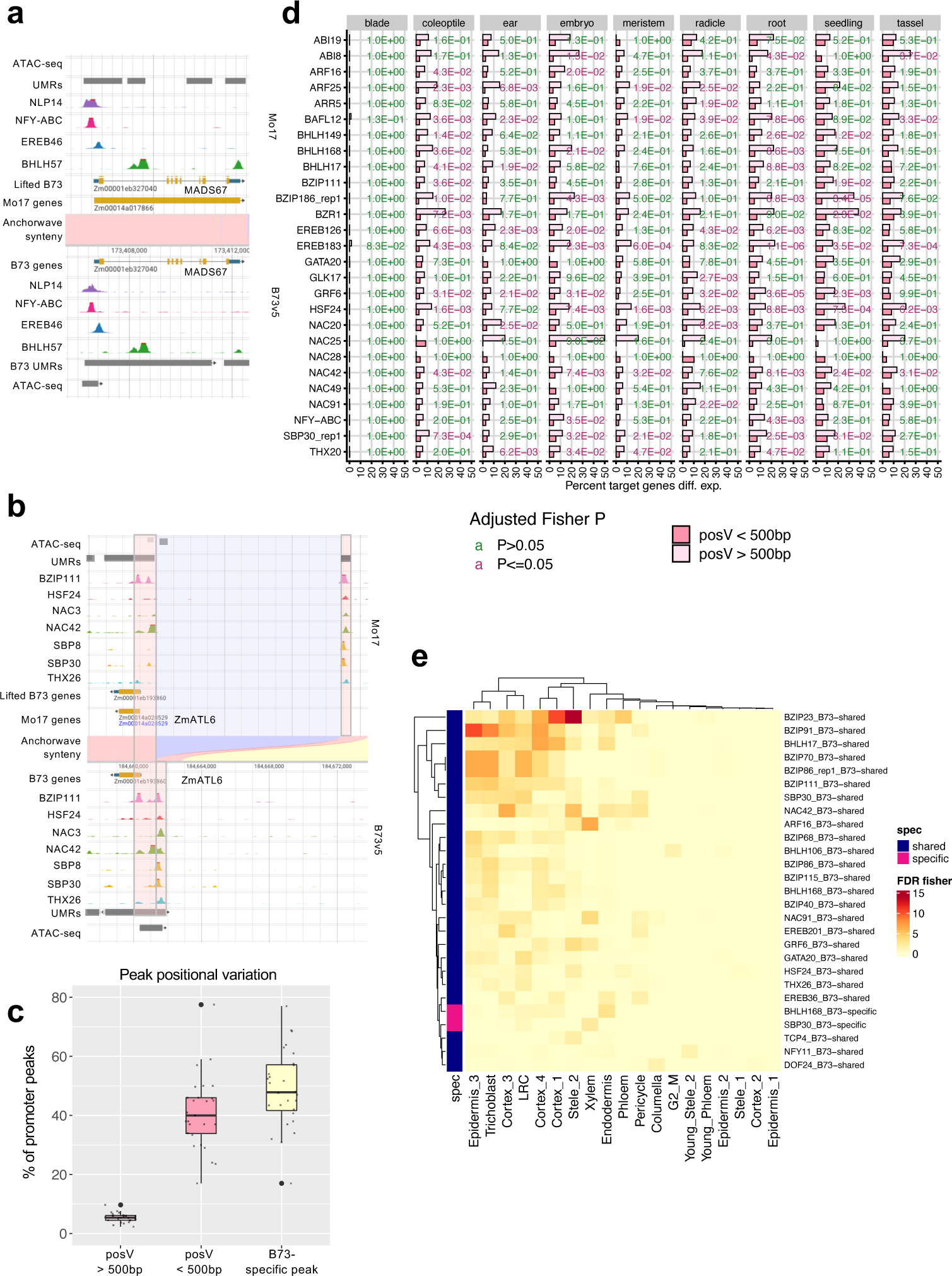
**a**. JBrowse 2 genome browser screenshot showing shared peaks of B73 and Mo17 at the *MADS67* locus. **b**. JBrowse 2 genome browser screenshot showing TF binding site positional variants at the *ZmATL6* locus. **c**. Percentage of B73 TF diversity panel promoter peaks that correspond to positional variants (posV) greater than 500bp, less than 500bp or B73-specific peaks overlapping structural variants. Each datapoint corresponds to the percentage of peaks from an individual TF overlapping the indicated category. **d.** Plot showing percentage of B73-Mo17 shared promoter peaks (< -10kb) that were associated with differentially expressed genes in various tissues from Zhou et al., 2019. Genes associated with posV<500 bp peaks are shown in dark pink, while those associated with posV>500 peaks are shown in light pink. Fisher’s exact tests were performed to assess the enrichment of DEGs in posV>500 bp genes relative to posV<500 bp genes for each TF and tissue combination, and the resulting P values were adjusted for multiple testing and shown. **e**. Heatmap showing the significance of association between putative target genes of shared or B73-specific peaks and root-type specific genes as determined by single cell RNA-seq data (Guillotin et al., 2023). B73-specific target genes are indicated with a magenta bar and shared target genes are indicated with a dark blue bar. Heatmap shows only a subset of TFs having an FDR <0.05 in at least one tissue. Heatmap color scale indicates FDR-adjusted Fisher’s pvalue.

**Supplemental Figure 13.**
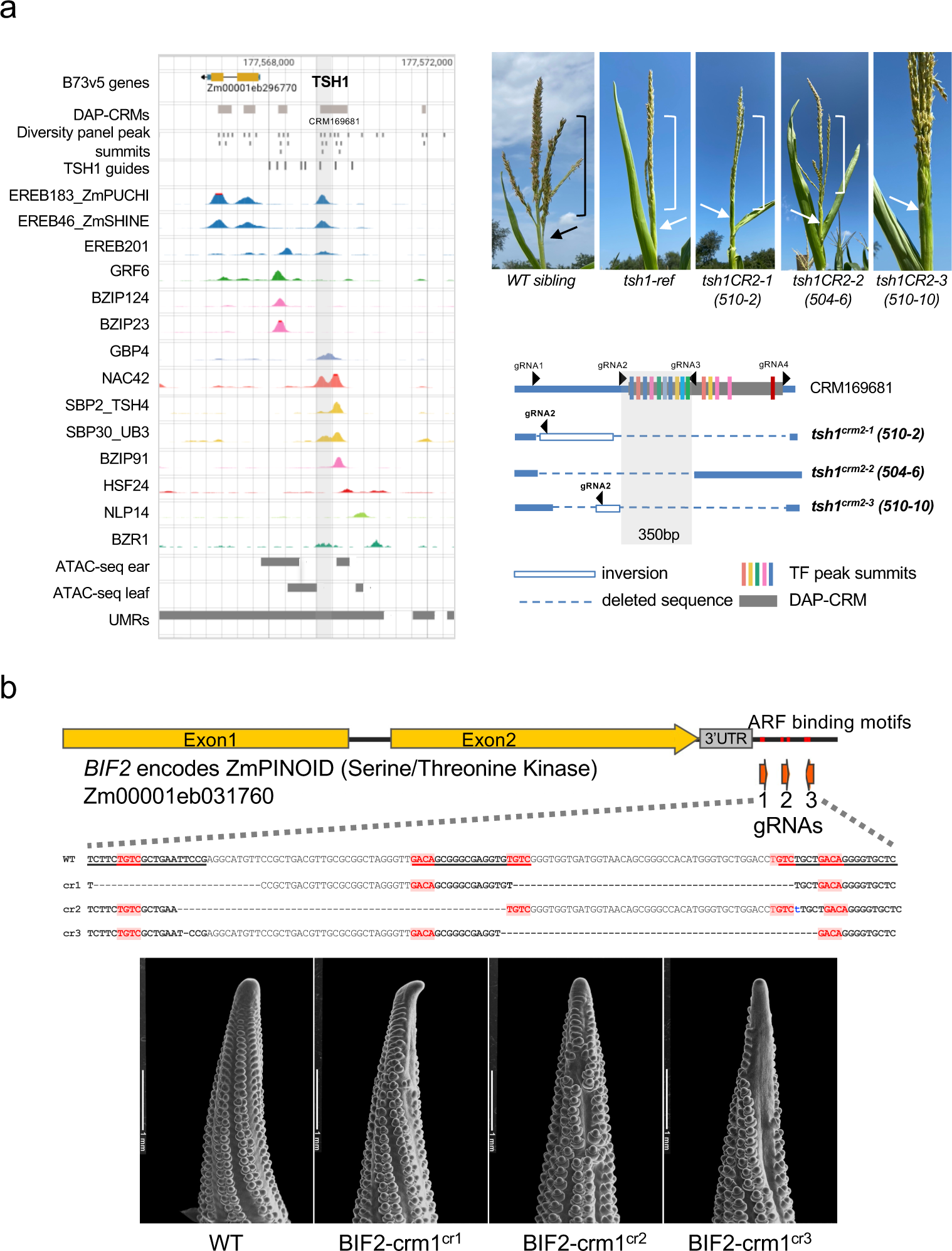
**a.** CRISPR editing of TSH1 regulatory regions. Left, genome browser screenshot of *TSH1* locus showing binding in upstream promoter region by many TFs. Grey shaded area within CRM169681 indicates region that was edited in alleles shown on lower right. Bottom right, schematic showing three independent CRISPR alleles with deletions and inversions that eliminate at least nine TF binding sites (colored bars; colors match TFs shown in genome browser screenshot). Lightly shaded grey area indicates a 350bp region that is common to all alleles and corresponds to the lightly shaded grey area in left panel. Top right, images of mature tassels for WT, *tsh1-ref* mutant (coding region mutation from Whipple et al. 2010), and the three TSH1 promoter CRISPR alleles shown below. All CRISPR promoter alleles show outgrowth of the tassel sheath leaf (white arrows) that is not present in the WT (black arrow). Tassel branching is reduced in the *tsh1-ref*, and the CRISPR promoter alleles (510-2 and 504-6; white brackets) relative to WT (black brackets). **b.** CRISPR editing of BIF2 3’ UTR ARF binding site. A *cis*-regulatory module (CRM18765) is situated downstream of the BIF2 gene. Within this CRM region, there is a strong ARF peak containing five ARF binding motifs (three TGTCs and two GACAs). Three single guide RNAs (gRNAs) were designed for CRISPR-Cas9 editing that specifically targeted the ARF motifs. Three deletion alleles were obtained. Homozygous plants of these alleles exhibited a weak *bif2* phenotype with various degrees of severity (*BIF2-crm1^cr1^* and *BIF2-crm1^cr3^*are more severe than *BIF2-crm1^cr2^*) during the early ear development stage, characterized by partial barren patches on the ear primordia.

**Supplemental Figure 14.**
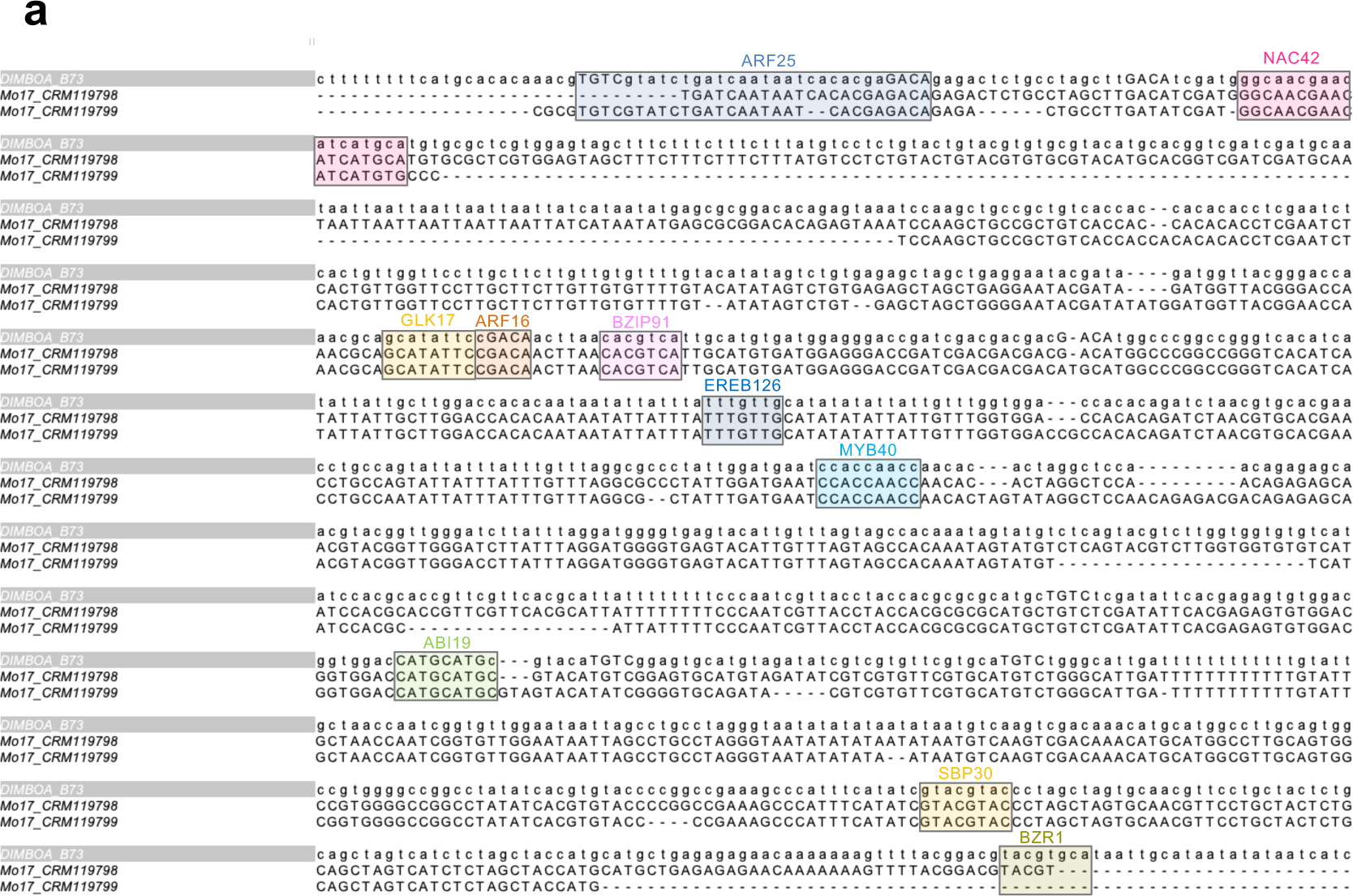
**a**. Nucleotide alignment of B73v3_DICE (single copy), Mo17_CRM119798 (tandem copy 1), and Mo17_CRM119799 (tandem copy 2) showing conservation of individual TF binding motifs.

## Notes

### Competing Interest Statement

The authors have declared no competing interest.

## REFERENCES

Aguirre, L., A. Hendelman, S. F. Hutton, D. M. McCandlish, and Z. B. Lippman. 2023. ’Idiosyncratic and dose-dependent epistasis drives variation in tomato fruit size’, Science, 382: 315–20.

Alonge, M., X. Wang, M. Benoit, S. Soyk, L. Pereira, L. Zhang, H. Suresh, S. Ramakrishnan, F. Maumus, D. Ciren, Y. Levy, T. H. Harel, G. Shalev-Schlosser, Z. Amsellem, H. Razifard, A. L. Caicedo, D. M. Tieman, H. Klee, M. Kirsche, S. Aganezov, T. R. Ranallo-Benavidez, Z. H. Lemmon, J. Kim, G. Robitaille, M. Kramer, S. Goodwin, W. R. McCombie, S. Hutton, J. Van Eck, J. Gillis, Y. Eshed, F. J. Sedlazeck, E. van der Knaap, M. C. Schatz, and Z. B. Lippman. 2020. ‘Major Impacts of Widespread Structural Variation on Gene Expression and Crop Improvement in Tomato’, Cell, 182: 145–61 e23.

Bang, S., X. Zhang, J. Gregory, Z. Chen, M. Galli, A. Gallavotti, and R.J. Schmitz. *WUSCHEL*-dependent chromatin regulation in maize inflorescence development at single-cell resolution bioRxiv 2024.05.13.593957

Barrera, L. A., A. Vedenko, J. V. Kurland, J. M. Rogers, S. S. Gisselbrecht, E. J. Rossin, J. Woodard, L. Mariani, K. H. Kock, S. Inukai, T. Siggers, L. Shokri, R. Gordan, N. Sahni, C. Cotsapas, T. Hao, S. Yi, M. Kellis, M. J. Daly, M. Vidal, D. E. Hill, and M. L. Bulyk. 2016. ‘Survey of variation in human transcription factors reveals prevalent DNA binding changes’, Science, 351: 1450–54.

Blanc-Mathieu, R., R. Dumas, L. Turchi, J. Lucas, and F. Parcy. 2024. ‘Plant-TFClass: a structural classification for plant transcription factors’, Trends Plant Sci, 29: 40–51.

Boer, D. R., A. Freire-Rios, W. A. van den Berg, T. Saaki, I. W. Manfield, S. Kepinski, I. Lopez-Vidrieo, J. M. Franco-Zorrilla, S. C. de Vries, R. Solano, D. Weijers, and M. Coll. 2014. ‘Structural basis for DNA binding specificity by the auxin-dependent ARF transcription factors’, Cell, 156: 577–89.

Bolger, A. M., M. Lohse, and B. Usadel. 2014. ‘Trimmomatic: a flexible trimmer for Illumina sequence data’, Bioinformatics, 30: 2114–20.

Buckler, E. S., J. B. Holland, P. J. Bradbury, C. B. Acharya, P. J. Brown, C. Browne, E. Ersoz, S. Flint-Garcia, A. Garcia, J. C. Glaubitz, M. M. Goodman, C. Harjes, K. Guill, D. E. Kroon, S. Larsson, N. K. Lepak, H. Li, S. E. Mitchell, G. Pressoir, J. A. Peiffer, M. O. Rosas, T. R. Rocheford, M. C. Romay, S. Romero, S. Salvo, H. Sanchez Villeda, H. S. da Silva, Q. Sun, F. Tian, N. Upadyayula, D. Ware, H. Yates, J. Yu, Z. Zhang, S. Kresovich, and M. D. McMullen. 2009. ‘The genetic architecture of maize flowering time’, Science, 325: 714–8.

Burdo, B., J. Gray, M. P. Goetting-Minesky, B. Wittler, M. Hunt, T. Li, D. Velliquette, J. Thomas, I. Gentzel, M. dos Santos Brito, M. K. Mejia-Guerra, L. N. Connolly, D. Qaisi, W. Li, M. I. Casas, A. I. Doseff, and E. Grotewold. 2014. ‘The Maize TFome--development of a transcription factor open reading frame collection for functional genomics’, Plant J, 80: 356–66.

Cahn, Jonathan, Michael Regulski, Jason Lynn, Evan Ernst, Cristiane de Santis Alves, Srividya Ramakrishnan, Kapeel Chougule, Sharon Wei, Zhenyuan Lu, Xiaosa Xu, Jorg Drenkow, Melissa Kramer, Arun Seetharam, Matthew B. Hufford, W. Richard McCombie, Doreen Ware, David Jackson, Michael C. Schatz, Thomas R. Gingeras, and Robert A. Martienssen. 2024. ‘MaizeCODE reveals bi-directionally expressed enhancers that harbor molecular signatures of maize domestication’, bioRxiv: 2024.02.22.581585.

Castelletti, S., R. Tuberosa, M. Pindo, and S. Salvi. 2014. ‘A MITE transposon insertion is associated with differential methylation at the maize flowering time QTL Vgt1’, G3 (Bethesda), 4: 805–12.

Chen, S., L. C. Francioli, J. K. Goodrich, R. L. Collins, M. Kanai, Q. Wang, J. Alfoldi, N. A. Watts, C. Vittal, L. D. Gauthier, T. Poterba, M. W. Wilson, Y. Tarasova, W. Phu, R. Grant, M. T. Yohannes, Z. Koenig, Y. Farjoun, E. Banks, S. Donnelly, S. Gabriel, N. Gupta, S. Ferriera, C. Tolonen, S. Novod, L. Bergelson, D. Roazen, V. Ruano-Rubio, M. Covarrubias, C. Llanwarne, N. Petrillo, G. Wade, T. Jeandet, R. Munshi, K. Tibbetts, Consortium Genome Aggregation Database, A. O’Donnell-Luria, M. Solomonson, C. Seed, A. R. Martin, M. E. Talkowski, H. L. Rehm, M. J. Daly, G. Tiao, B. M. Neale, D. G. MacArthur, and K. J. Karczewski. 2024. ’A genomic mutational constraint map using variation in 76,156 human genomes’, Nature, 625: 92–100.

Chuck, G. S., P. J. Brown, R. Meeley, and S. Hake. 2014. ‘Maize SBP-box transcription factors unbranched2 and unbranched3 affect yield traits by regulating the rate of lateral primordia initiation’, Proc Natl Acad Sci U S A, 111: 18775–80.

Crawford, B. C., and M. F. Yanofsky. 2011. ‘HALF FILLED promotes reproductive tract development and fertilization efficiency in Arabidopsis thaliana’, Development, 138: 2999–3009.

Crisp, P. A., A. P. Marand, J. M. Noshay, P. Zhou, Z. Lu, R. J. Schmitz, and N. M. Springer. 2020. ‘Stable unmethylated DNA demarcates expressed genes and their cis-regulatory space in plant genomes’, Proc Natl Acad Sci U S A, 117: 23991–4000.

Dai, D., J. S. Mudunkothge, M. Galli, S. N. Char, R. Davenport, X. Zhou, J. L. Gustin, G. Spielbauer, J. Zhang, W. B. Barbazuk, B. Yang, A. Gallavotti, and A. M. Settles. 2022. ‘Paternal imprinting of dosage-effect defective1 contributes to seed weight xenia in maize’, Nat Commun, 13: 5366.

de Martin, X., R. Sodaei, and G. Santpere. 2021. ‘Mechanisms of Binding Specificity among bHLH Transcription Factors’, Int J Mol Sci, 22.

De Masi, F., C. A. Grove, A. Vedenko, A. Alibes, S. S. Gisselbrecht, L. Serrano, M. L. Bulyk, and A. J. Walhout. 2011. ‘Using a structural and logics systems approach to infer bHLH-DNA binding specificity determinants’, Nucleic Acids Res, 39: 4553–63.

Diesh, C., G. J. Stevens, P. Xie, T. De Jesus Martinez, E. A. Hershberg, A. Leung, E. Guo, S. Dider, J. Zhang, C. Bridge, G. Hogue, A. Duncan, M. Morgan, T. Flores, B. N. Bimber, R. Haw, S. Cain, R. M. Buels, L. D. Stein, and I. H. Holmes. 2023. ‘JBrowse 2: a modular genome browser with views of synteny and structural variation’, Genome Biol, 24: 74.

Dobin, A., C. A. Davis, F. Schlesinger, J. Drenkow, C. Zaleski, S. Jha, P. Batut, M. Chaisson, and T. R. Gingeras. 2013. ‘STAR: ultrafast universal RNA-seq aligner’, Bioinformatics, 29: 15–21.

Dong, Z., Z. Xu, L. Xu, M. Galli, A. Gallavotti, H. K. Dooner, and G. Chuck. 2020. ‘Necrotic upper tips1 mimics heat and drought stress and encodes a protoxylem-specific transcription factor in maize’, Proc Natl Acad Sci U S A, 117: 20908–19.

Eichten, S. R., J. M. Foerster, N. de Leon, Y. Kai, C. T. Yeh, S. Liu, J. A. Jeddeloh, P. S. Schnable, S. M. Kaeppler, and N. M. Springer. 2011. ‘B73-Mo17 near-isogenic lines demonstrate dispersed structural variation in maize’, Plant Physiol, 156: 1679–90.

Engelhorn, Julia, Samantha J. Snodgrass, Amelie Kok, Arun S. Seetharam, Michael Schneider, Tatjana Kiwit, Ayush Singh, Michael Banf, Merritt Khaipho-Burch, Daniel E. Runcie, Victor Sánchez Camargo, J. Vladimir Torres-Rodriguez, Guangchao Sun, Maike Stam, Fabio Fiorani, James C. Schnable, Hank W. Bass, Matthew B. Hufford, Benjamin Stich, Wolf B. Frommer, Jeffrey Ross-Ibarra, and Thomas Hartwig. 2023. ‘Phenotypic variation in maize can be largely explained by genetic variation at transcription factor binding sites’, bioRxiv: 2023.08.08.551183.

Evans, M. M. 2007. ‘The indeterminate gametophyte1 gene of maize encodes a LOB domain protein required for embryo Sac and leaf development’, Plant Cell, 19: 46–62.

Flatschacher, D., V. Speckbacher, and S. Zeilinger. 2022. ‘qRAT: an R-based stand-alone application for relative expression analysis of RT-qPCR data’, BMC Bioinformatics, 23: 286.

Frey, M., P. Chomet, E. Glawischnig, C. Stettner, S. Grun, A. Winklmair, W. Eisenreich, A. Bacher, R. B. Meeley, S. P. Briggs, K. Simcox, and A. Gierl. 1997. ‘Analysis of a chemical plant defense mechanism in grasses’, Science, 277: 696–9.

Gaillochet, C., T. Stiehl, C. Wenzl, J. J. Ripoll, L. J. Bailey-Steinitz, L. Li, A. Pfeiffer, A. Miotk, J. P. Hakenjos, J. Forner, M. F. Yanofsky, A. Marciniak-Czochra, and J. U. Lohmann. 2017. ‘Control of plant cell fate transitions by transcriptional and hormonal signals’, Elife, 6.

Gallavotti, A., Q. Zhao, J. Kyozuka, R. B. Meeley, M. K. Ritter, J. F. Doebley, M. E. Pe, and R. J. Schmidt. 2004. ‘The role of barren stalk1 in the architecture of maize’, Nature, 432: 630–5.

Galli, M., A. Khakhar, Z. Lu, Z. Chen, S. Sen, T. Joshi, J. L. Nemhauser, R. J. Schmitz, and A. Gallavotti. 2018. ‘The DNA binding landscape of the maize AUXIN RESPONSE FACTOR family’, Nat Commun, 9: 4526.

Ge, S. X., D. Jung, and R. Yao. 2020. ‘ShinyGO: a graphical gene-set enrichment tool for animals and plants’, Bioinformatics, 36: 2628–29.

Girin, T., T. Paicu, P. Stephenson, S. Fuentes, E. Korner, M. O’Brien, K. Sorefan, T. A. Wood, V. Balanza, C. Ferrandiz, D. R. Smyth, and L. Ostergaard. 2011. ‘INDEHISCENT and SPATULA interact to specify carpel and valve margin tissue and thus promote seed dispersal in Arabidopsis’, Plant Cell, 23: 3641–53.

Goel, M., H. Sun, W. B. Jiao, and K. Schneeberger. 2019. ‘SyRI: finding genomic rearrangements and local sequence differences from whole-genome assemblies’, Genome Biol, 20: 277.

Gremski, K., G. Ditta, and M. F. Yanofsky. 2007. ‘The HECATE genes regulate female reproductive tract development in Arabidopsis thaliana’, Development, 134: 3593–601.

Gu, Z., R. Eils, and M. Schlesner. 2016. ‘Complex heatmaps reveal patterns and correlations in multidimensional genomic data’, Bioinformatics, 32: 2847–9.

Gu, Zuguang. 2022. ‘Complex heatmap visualization’, iMeta, 1: e43.

Guillotin, B., R. Rahni, M. Passalacqua, M. A. Mohammed, X. Xu, S. K. Raju, C. O. Ramirez, D. Jackson, S. C. Groen, J. Gillis, and K. D. Birnbaum. 2023. ‘A pan-grass transcriptome reveals patterns of cellular divergence in crops’, Nature, 617: 785–91.

Guo, Y., S. Mahony, and D. K. Gifford. 2012. ‘High resolution genome wide binding event finding and motif discovery reveals transcription factor spatial binding constraints’, PLoS Comput Biol, 8: e1002638.

Hajheidari, M., and S. C. Huang. 2022. ‘Elucidating the biology of transcription factor-DNA interaction for accurate identification of cis-regulatory elements’, Curr Opin Plant Biol, 68: 102232.

Heim, M. A., M. Jakoby, M. Werber, C. Martin, B. Weisshaar, and P. C. Bailey. 2003. ‘The basic helix-loop-helix transcription factor family in plants: a genome-wide study of protein structure and functional diversity’, Mol Biol Evol, 20: 735–47.

Hermann, M., F. Maier, A. Masroor, S. Hirth, A. J. Pfitzner, and U. M. Pfitzner. 2013. ‘The Arabidopsis NIMIN proteins affect NPR1 differentially’, Front Plant Sci, 4: 88.

Hufford, M. B., A. S. Seetharam, M. R. Woodhouse, K. M. Chougule, S. Ou, J. Liu, W. A. Ricci, T. Guo, A. Olson, Y. Qiu, R. Della Coletta, S. Tittes, A. I. Hudson, A. P. Marand, S. Wei, Z. Lu, B. Wang, M. K. Tello-Ruiz, R. D. Piri, N. Wang, D. W. Kim, Y. Zeng, C. H. O’Connor, X. Li, A. M. Gilbert, E. Baggs, K. V. Krasileva, J. L. Portwood, 2nd, E. K. S. Cannon, C. M. Andorf, N. Manchanda, S. J. Snodgrass, D. E. Hufnagel, Q. Jiang, S. Pedersen, M. L. Syring, D. A. Kudrna, V. Llaca, K. Fengler, R. J. Schmitz, J. Ross-Ibarra, J. Yu, J. I. Gent, C. N. Hirsch, D. Ware, and R. K. Dawe. 2021. ‘De novo assembly, annotation, and comparative analysis of 26 diverse maize genomes’, Science, 373: 655–62.

Hugouvieux, V., and C. Zubieta. 2018. ‘MADS transcription factors cooperate: complexities of complex formation’, J Exp Bot, 69: 1821–23.

Iotchkova, V., G. R. S. Ritchie, M. Geihs, S. Morganella, J. L. Min, K. Walter, N. J. Timpson, Uk K. Consortium, I. Dunham, E. Birney, and N. Soranzo. 2019. ‘GARFIELD classifies disease-relevant genomic features through integration of functional annotations with association signals’, Nat Genet, 51: 343–53.

Jores, T., J. Tonnies, N. A. Mueth, A. Romanowski, S. Fields, J. T. Cuperus, and C. Queitsch. 2024. ‘Plant enhancers exhibit both cooperative and additive interactions among their functional elements’, Plant Cell.

Jumper, J., R. Evans, A. Pritzel, T. Green, M. Figurnov, O. Ronneberger, K. Tunyasuvunakool, R. Bates, A. Zidek, A. Potapenko, A. Bridgland, C. Meyer, S. A. A. Kohl, A. J. Ballard, A. Cowie, B. Romera-Paredes, S. Nikolov, R. Jain, J. Adler, T. Back, S. Petersen, D. Reiman, E. Clancy, M. Zielinski, M. Steinegger, M. Pacholska, T. Berghammer, D. Silver, O. Vinyals, A. W. Senior, K. Kavukcuoglu, P. Kohli, and D. Hassabis. 2021. ‘Applying and improving AlphaFold at CASP14’, Proteins, 89: 1711–21.

Kielbasa, S. M., R. Wan, K. Sato, P. Horton, and M. C. Frith. 2011. ‘Adaptive seeds tame genomic sequence comparison’, Genome Res, 21: 487–93.

Kim, S., and J. Wysocka. 2023. ‘Deciphering the multi-scale, quantitative cis-regulatory code’, Mol Cell, 83: 373–92.

Lai, X., A. Stigliani, J. Lucas, V. Hugouvieux, F. Parcy, and C. Zubieta. 2020. ‘Genome-wide binding of SEPALLATA3 and AGAMOUS complexes determined by sequential DNA-affinity purification sequencing’, Nucleic Acids Res, 48: 9637–48.

Langmead, B., and S. L. Salzberg. 2012. ‘Fast gapped-read alignment with Bowtie 2’, Nat Methods, 9: 357–9.

Li, B., and C. N. Dewey. 2011. ‘RSEM: accurate transcript quantification from RNA-Seq data with or without a reference genome’, BMC Bioinformatics, 12: 323.

Li, M., T. Yao, W. Lin, W. E. Hinckley, M. Galli, W. Muchero, A. Gallavotti, J. G. Chen, and S. C. Huang. 2023. ‘Double DAP-seq uncovered synergistic DNA binding of interacting bZIP transcription factors’, Nat Commun, 14: 2600.

Liang, Y., Q. Liu, X. Wang, C. Huang, G. Xu, S. Hey, H. Y. Lin, C. Li, D. Xu, L. Wu, C. Wang, W. Wu, J. Xia, X. Han, S. Lu, J. Lai, W. Song, P. S. Schnable, and F. Tian. 2019. ‘ZmMADS69 functions as a flowering activator through the ZmRap2.7-ZCN8 regulatory module and contributes to maize flowering time adaptation’, New Phytol, 221: 2335–47.

Lu, Z., A. P. Marand, W. A. Ricci, C. L. Ethridge, X. Zhang, and R. J. Schmitz. 2019. ‘The prevalence, evolution and chromatin signatures of plant regulatory elements’, Nat Plants, 5: 1250–59.

Machanick, P., and T. L. Bailey. 2011. ‘MEME-ChIP: motif analysis of large DNA datasets’, Bioinformatics, 27: 1696–7.

Maekawa, S., T. Sato, Y. Asada, S. Yasuda, M. Yoshida, Y. Chiba, and J. Yamaguchi. 2012. ‘The Arabidopsis ubiquitin ligases ATL31 and ATL6 control the defense response as well as the carbon/nitrogen response’, Plant Mol Biol, 79: 217–27.

Majer, C., and F. Hochholdinger. 2011. ‘Defining the boundaries: structure and function of LOB domain proteins’, Trends Plant Sci, 16: 47–52.

Marand, A. P., A. L. Eveland, K. Kaufmann, and N. M. Springer. 2023. ‘cis-Regulatory Elements in Plant Development, Adaptation, and Evolution’, Annu Rev Plant Biol, 74: 111–37.

Maurano, M. T., R. Humbert, E. Rynes, R. E. Thurman, E. Haugen, H. Wang, A. P. Reynolds, R. Sandstrom, H. Qu, J. Brody, A. Shafer, F. Neri, K. Lee, T. Kutyavin, S. Stehling-Sun, A. K. Johnson, T. K. Canfield, E. Giste, M. Diegel, D. Bates, R. S. Hansen, S. Neph, P. J. Sabo, S. Heimfeld, A. Raubitschek, S. Ziegler, C. Cotsapas, N. Sotoodehnia, I. Glass, S. R. Sunyaev, R. Kaul, and J. A. Stamatoyannopoulos. 2012. ‘Systematic localization of common disease-associated variation in regulatory DNA’, Science, 337: 1190–5.

McSteen, P., S. Malcomber, A. Skirpan, C. Lunde, X. Wu, E. Kellogg, and S. Hake. 2007. ‘barren inflorescence2 Encodes a co-ortholog of the PINOID serine/threonine kinase and is required for organogenesis during inflorescence and vegetative development in maize’, Plant Physiol, 144: 1000–11.

Mirdita, M., K. Schutze, Y. Moriwaki, L. Heo, S. Ovchinnikov, and M. Steinegger. 2022. ‘ColabFold: making protein folding accessible to all’, Nat Methods, 19: 679–82.

Noshay, J. M., Z. Liang, P. Zhou, P. A. Crisp, A. P. Marand, C. N. Hirsch, R. J. Schmitz, and N. M. Springer. 2021. ‘Stability of DNA methylation and chromatin accessibility in structurally diverse maize genomes’, G3 (Bethesda), 11.

O’Malley, R. C., S. C. Huang, L. Song, M. G. Lewsey, A. Bartlett, J. R. Nery, M. Galli, A. Gallavotti, and J. R. Ecker. 2016. ’Cistrome and Epicistrome Features Shape the Regulatory DNA Landscape’, Cell, 166: 1598.

Oka, R., J. Zicola, B. Weber, S. N. Anderson, C. Hodgman, J. I. Gent, J. J. Wesselink, N. M. Springer, H. C. J. Hoefsloot, F. Turck, and M. Stam. 2017. ‘Genome-wide mapping of transcriptional enhancer candidates using DNA and chromatin features in maize’, Genome Biol, 18: 137.

Parvathaneni, R. K., E. Bertolini, M. Shamimuzzaman, D. L. Vera, P. Y. Lung, B. R. Rice, J. Zhang, P. J. Brown, A. E. Lipka, H. W. Bass, and A. L. Eveland. 2020. ‘The regulatory landscape of early maize inflorescence development’, Genome Biol, 21: 165.

Peng, B., X. Zhao, Y. Wang, C. Li, Y. Li, D. Zhang, Y. Shi, Y. Song, L. Wang, Y. Li, and T. Wang. 2021. ‘Genome-wide association studies of leaf angle in maize’, Mol Breed, 41: 50.

Quinlan, A. R., and I. M. Hall. 2010. ‘BEDTools: a flexible suite of utilities for comparing genomic features’, Bioinformatics, 26: 841–2.

Ramirez, F., F. Dundar, S. Diehl, B. A. Gruning, and T. Manke. 2014. ‘deepTools: a flexible platform for exploring deep-sequencing data’, Nucleic Acids Res, 42: W187–91.

Ricci, W. A., Z. Lu, L. Ji, A. P. Marand, C. L. Ethridge, N. G. Murphy, J. M. Noshay, M. Galli, M. K. Mejia-Guerra, M. Colome-Tatche, F. Johannes, M. J. Rowley, V. G. Corces, J. Zhai, M. J. Scanlon, E. S. Buckler, A. Gallavotti, N. M. Springer, R. J. Schmitz, and X. Zhang. 2019. ‘Widespread long-range cis-regulatory elements in the maize genome’, Nat Plants, 5: 1237–49.

Richardson, A. E., and S. Hake. 2022. ‘The power of classic maize mutants: Driving forward our fundamental understanding of plants’, Plant Cell, 34: 2505–17.

Ritchie, M. E., B. Phipson, D. Wu, Y. Hu, C. W. Law, W. Shi, and G. K. Smyth. 2015. ‘limma powers differential expression analyses for RNA-sequencing and microarray studies’, Nucleic Acids Res, 43: e47.

Robinson, J. T., H. Thorvaldsdottir, W. Winckler, M. Guttman, E. S. Lander, G. Getz, and J. P. Mesirov. 2011. ‘Integrative genomics viewer’, Nat Biotechnol, 29: 24–6.

Rodgers-Melnick, E., D. L. Vera, H. W. Bass, and E. S. Buckler. 2016. ‘Open chromatin reveals the functional maize genome’, Proc Natl Acad Sci U S A, 113: E3177–84.

Rodriguez-Leal, D., Z. H. Lemmon, J. Man, M. E. Bartlett, and Z. B. Lippman. 2017. ‘Engineering Quantitative Trait Variation for Crop Improvement by Genome Editing’, Cell, 171: 470–80 e8.

Salvi, S., G. Sponza, M. Morgante, D. Tomes, X. Niu, K. A. Fengler, R. Meeley, E. V. Ananiev, S. Svitashev, E. Bruggemann, B. Li, C. F. Hainey, S. Radovic, G. Zaina, J. A. Rafalski, S. V. Tingey, G. H. Miao, R. L. Phillips, and R. Tuberosa. 2007. ‘Conserved noncoding genomic sequences associated with a flowering-time quantitative trait locus in maize’, Proc Natl Acad Sci U S A, 104: 11376–81.

Savadel, S. D., T. Hartwig, Z. M. Turpin, D. L. Vera, P. Y. Lung, X. Sui, M. Blank, W. B. Frommer, J. H. Dennis, J. Zhang, and H. W. Bass. 2021. ‘The native cistrome and sequence motif families of the maize ear’, PLoS Genet, 17: e1009689.

Schmitz, R. J., E. Grotewold, and M. Stam. 2022. ‘Cis-regulatory sequences in plants: Their importance, discovery, and future challenges’, Plant Cell, 34: 718–41.

Shen, Z., M. A. Hoeksema, Z. Ouyang, C. Benner, and C. K. Glass. 2020. ‘MAGGIE: leveraging genetic variation to identify DNA sequence motifs mediating transcription factor binding and function’, Bioinformatics, 36: i84–i92.

Shumate, A., and S. L. Salzberg. 2021. ‘Liftoff: accurate mapping of gene annotations’, Bioinformatics, 37: 1639–43.

Smaczniak, C., J. M. Muino, D. Chen, G. C. Angenent, and K. Kaufmann. 2017. ’Differences in DNA Binding Specificity of Floral Homeotic Protein Complexes Predict Organ-Specific Target Genes’, *Plant Cell*, 29: 1822-35. Soneson, C., M. I. Love, and M. D. Robinson. 2015. ‘Differential analyses for RNA-seq: transcript-level estimates improve gene-level inferences’, F1000Res, 4: 1521.

Song, B., E. S. Buckler, H. Wang, Y. Wu, E. Rees, E. A. Kellogg, D. J. Gates, M. Khaipho-Burch, P. J. Bradbury, J. Ross-Ibarra, M. B. Hufford, and M. C. Romay. 2021. ‘Conserved noncoding sequences provide insights into regulatory sequence and loss of gene expression in maize’, Genome Res, 31: 1245–57.

Song, B., S. Marco-Sola, M. Moreto, L. Johnson, E. S. Buckler, and M. C. Stitzer. 2022. ‘AnchorWave: Sensitive alignment of genomes with high sequence diversity, extensive structural polymorphism, and whole-genome duplication’, Proc Natl Acad Sci U S A, 119.

Spielmann, M., and S. Mundlos. 2016. ‘Looking beyond the genes: the role of non-coding variants in human disease’, Hum Mol Genet, 25: R157–R65.

Springer, N., N. de Leon, and E. Grotewold. 2019. ‘Challenges of Translating Gene Regulatory Information into Agronomic Improvements’, Trends Plant Sci, 24: 1075–82.

Sun, S., Y. Zhou, J. Chen, J. Shi, H. Zhao, H. Zhao, W. Song, M. Zhang, Y. Cui, X. Dong, H. Liu, X. Ma, Y. Jiao, B. Wang, X. Wei, J. C. Stein, J. C. Glaubitz, F. Lu, G. Yu, C. Liang, K. Fengler, B. Li, A. Rafalski, P. S. Schnable, D. H. Ware, E. S. Buckler, and J. Lai. 2018. ‘Extensive intraspecific gene order and gene structural variations between Mo17 and other maize genomes’, Nat Genet, 50: 1289–95.

Sun, Y., L. Dong, Y. Zhang, D. Lin, W. Xu, C. Ke, L. Han, L. Deng, G. Li, D. Jackson, X. Li, and F. Yang. 2020. ‘3D genome architecture coordinates trans and cis regulation of differentially expressed ear and tassel genes in maize’, Genome Biol, 21: 143.

Tu, X., M. K. Mejia-Guerra, J. A. Valdes Franco, D. Tzeng, P. Y. Chu, W. Shen, Y. Wei, X. Dai, P. Li, E. S. Buckler, and S. Zhong. 2020. ‘Reconstructing the maize leaf regulatory network using ChIP-seq data of 104 transcription factors’, Nat Commun, 11: 5089.

Wallace, J. G., P. J. Bradbury, N. Zhang, Y. Gibon, M. Stitt, and E. S. Buckler. 2014. ‘Association mapping across numerous traits reveals patterns of functional variation in maize’, PLoS Genet, 10: e1004845.

Wallace, J. G., E. Rodgers-Melnick, and E. S. Buckler. 2018. ‘On the Road to Breeding 4.0: Unraveling the Good, the Bad, and the Boring of Crop Quantitative Genomics’, Annu Rev Genet, 52: 421–44.

Weirauch, M. T., A. Yang, M. Albu, A. G. Cote, A. Montenegro-Montero, P. Drewe, H. S. Najafabadi, S. A. Lambert, I. Mann, K. Cook, H. Zheng, A. Goity, H. van Bakel, J. C. Lozano, M. Galli, M. G. Lewsey, E. Huang, T. Mukherjee, X. Chen, J. S. Reece-Hoyes, S. Govindarajan, G. Shaulsky, A. J. M. Walhout, F. Y. Bouget, G. Ratsch, L. F. Larrondo, J. R. Ecker, and T. R. Hughes. 2014. ‘Determination and inference of eukaryotic transcription factor sequence specificity’, Cell, 158: 1431–43.

Whipple, C. J., D. H. Hall, S. DeBlasio, F. Taguchi-Shiobara, R. J. Schmidt, and D. P. Jackson. 2010. ‘A conserved mechanism of bract suppression in the grass family’, Plant Cell, 22: 565–78.

Wu, H., M. Galli, C. J. Spears, J. Zhan, P. Liu, R. Yadegari, J. M. Dannenhoffer, A. Gallavotti, and P. W. Becraft. 2023. ‘NAKED ENDOSPERM1, NAKED ENDOSPERM2, and OPAQUE2 interact to regulate gene networks in maize endosperm development’, Plant Cell, 36: 19–39.

Wu, J., S. J. Lawit, B. Weers, J. Sun, N. Mongar, J. Van Hemert, R. Melo, X. Meng, M. Rupe, J. Clapp, K. Haug Collet, L. Trecker, K. Roesler, L. Peddicord, J. Thomas, J. Hunt, W. Zhou, Z. Hou, M. Wimmer, J. Jantes, H. Mo, L. Liu, Y. Wang, C. Walker, O. Danilevskaya, R. H. Lafitte, J. R. Schussler, B. Shen, and J. E. Habben. 2019. ‘Overexpression of zmm28 increases maize grain yield in the field’, Proc Natl Acad Sci U S A, 116: 23850–58.

Xiao, Y., J. Guo, Z. Dong, A. Richardson, E. Patterson, S. Mangrum, S. Bybee, E. Bertolini, M. Bartlett, G. Chuck, A. L. Eveland, M. J. Scanlon, and C. Whipple. 2022. ’Boundary domain genes were recruited to suppress bract growth and promote branching in maize’, Sci Adv, 8: eabm6835.

Xing, H. L., L. Dong, Z. P. Wang, H. Y. Zhang, C. Y. Han, B. Liu, X. C. Wang, and Q. J. Chen. 2014. ‘A CRISPR/Cas9 toolkit for multiplex genome editing in plants’, BMC Plant Biol, 14: 327.

Xu, C., H. Cao, Q. Zhang, H. Wang, W. Xin, E. Xu, S. Zhang, R. Yu, D. Yu, and Y. Hu. 2018. ‘Control of auxin-induced callus formation by bZIP59-LBD complex in Arabidopsis regeneration’, Nat Plants, 4: 108–15.

Yilmaz, A., M. Y. Nishiyama, Jr., B. G. Fuentes, G. M. Souza, D. Janies, J. Gray, and E. Grotewold. 2009. ‘GRASSIUS: a platform for comparative regulatory genomics across the grasses’, Plant Physiol, 149: 171–80.

Yocca, A. E., and P. P. Edger. 2022. ‘Current status and future perspectives on the evolution of cis-regulatory elements in plants’, Curr Opin Plant Biol, 65: 102139.

Yu, G., L. G. Wang, and Q. Y. He. 2015. ‘ChIPseeker: an R/Bioconductor package for ChIP peak annotation, comparison and visualization’, Bioinformatics, 31: 2382–3.

Zhan, J., G. Li, C. H. Ryu, C. Ma, S. Zhang, A. Lloyd, B. G. Hunter, B. A. Larkins, G. N. Drews, X. Wang, and R. Yadegari. 2018. ‘Opaque-2 Regulates a Complex Gene Network Associated with Cell Differentiation and Storage Functions of Maize Endosperm’, Plant Cell, 30: 2425–46.

Zhang, Y., Z. Li, Y. Zhang, K. Lin, Y. Peng, L. Ye, Y. Zhuang, M. Wang, Y. Xie, J. Guo, W. Teng, Y. Tong, W. Zhang, Y. Xue, Z. Lang, and Y. Zhang. 2021. ‘Evolutionary rewiring of the wheat transcriptional regulatory network by lineage-specific transposable elements’, Genome Res, 31: 2276–89.

Zhao, H., Z. Sun, J. Wang, H. Huang, J. P. Kocher, and L. Wang. 2014. ‘CrossMap: a versatile tool for coordinate conversion between genome assemblies’, Bioinformatics, 30: 1006–7.

Zheng, L., M. D. McMullen, E. Bauer, C. C. Schon, A. Gierl, and M. Frey. 2015. ‘Prolonged expression of the BX1 signature enzyme is associated with a recombination hotspot in the benzoxazinoid gene cluster in Zea mays’, J Exp Bot, 66: 3917–30.

Zhou, P., C. N. Hirsch, S. P. Briggs, and N. M. Springer. 2019. ‘Dynamic Patterns of Gene Expression Additivity and Regulatory Variation throughout Maize Development’, Mol Plant, 12: 410–25.

